# Modulation of sensory perception by hydrogen peroxide enables Caenorhabditis elegans to find a niche that provides both food and protection from hydrogen peroxide

**DOI:** 10.1101/2021.09.23.461430

**Authors:** Jodie A. Schiffer, Stephanie V. Stumbur, Maedeh Seyedolmohadesin, Yuyan Xu, William T. Serkin, Natalie G McGowan, Oluwatosin Banjo, Mahdi Torkashvand, Albert Lin, Ciara N. Hosea, Adrien Assié, Buck S. Samuel, Michael P. O’Donnell, Vivek Venkatachalam, Javier Apfeld

## Abstract

Hydrogen peroxide (H_2_O_2_) is the most common chemical threat that organisms face. Here, we show that H_2_O_2_ alters the bacterial food preference of *Caenorhabditis elegans*, enabling the nematodes to find a safe environment with food. H_2_O_2_ induces the nematodes to leave food patches of laboratory and microbiome bacteria when those bacterial communities have insufficient H_2_O_2_-degrading capacity. The nematode’s behavior is directed by H_2_O_2_-sensing neurons that promote escape from H_2_O_2_ and by bacteria-sensing neurons that promote attraction to bacteria. However, the input for H_2_O_2_-sensing neurons is removed by bacterial H_2_O_2_-degrading enzymes and the bacteria-sensing neurons’ perception of bacteria is prevented by H_2_O_2_. The resulting cross-attenuation provides a general mechanism that ensures the nematode’s behavior is faithful to the lethal threat of hydrogen peroxide, increasing the nematode’s chances of finding a niche that provides both food and protection from hydrogen peroxide.

## Introduction

To grow and reproduce in an ever-changing natural environment, animals must adjust their behavior to find both food and safety. Animals co-evolved in close association with complex bacterial communities that can remodel both the animals’ behavior and their environment (Ezenwa et al., 2012; McFall-Ngai et al., 2013). Because of this complexity, our understanding of the evolution of the mechanisms that adjust animal behavior to enable them to find food and safety in variable environments remains limited (Gordon, 2016). In the present study, we developed a model ecosystem to determine how the environment-dependent sensory perception of the natural bacterial community enables *Caenorhabditis elegans* nematodes to adjust their behavior and find a niche that provides both food and protection from hydrogen peroxide.

Hydrogen peroxide (H_2_O_2_) is the most common chemical threat in the microbial battlefield (Mishra and Imlay, 2012). Bacteria, fungi, plants, and animal cells have long been known to excrete H_2_O_2_ to attack prey and pathogens (Avery and Morgan, 1924; Imlay, 2018). H_2_O_2_ is also a byproduct of aerobic respiration (Chance et al., 1979). Prevention and repair of the damage that hydrogen peroxide inflicts on macromolecules are critical for cellular health and survival (Chance et al., 1979). To avoid damage from H_2_O_2_, cells rely on conserved physiological defenses, including H_2_O_2_-degrading catalases (Mishra and Imlay, 2012). We recently found that *C. elegans* represses their own H_2_O_2_ defenses in response to sensory perception of *Escherichia coli*, the nematode’s food source, because *E. coli* can deplete H_2_O_2_ from the local environment and thereby protect the nematodes (Schiffer et al., 2020). Thus, the *E. coli* self-defense mechanisms create a public good (West et al., 2006), an environment safe from the threat of H_2_O_2_, that benefits *C. elegans* (Schiffer et al., 2020). Whether similar interactions between nematodes and bacteria shaped the evolution of behavioral responses protecting *C. elegans* from H_2_O_2_ remains poorly understood.

*C. elegans* is ideally suited for studying how bacteria shape the evolution of behaviors that enable animals to find food and H_2_O_2_ protection because of *C. elegans’* small size, well-described anatomy (Cook et al., 2019; Sulston et al., 1983), and tractable microbiome (Berg et al., 2016; Dirksen et al., 2020; Dirksen et al., 2016; O’Donnell et al., 2020; Samuel et al., 2016). *C. elegans* associates with a bacterial microbiome recruited from the surrounding environment (Berg et al., 2016; Dirksen et al., 2016; Samuel et al., 2016) that includes bacteria in genera that degrade or produce H_2_O_2_ (Passardi et al., 2007). Hydrogen peroxide produced by a bacterium from the *C. elegans* microbiome, *Neorhizobium sp*., causes DNA damage to the nematodes (Kniazeva and Ruvkun, 2019). Many bacteria—including *Streptococcus pyogenes, Streptococcus pneumoniae, Streptococcus oralis*, and *Enterococcus faecium*—kill *C. elegans* by producing millimolar concentrations of H_2_O_2_ (Bolm et al., 2004; Jansen et al., 2002; Moy et al., 2004). In the complex and variable habitat where *C. elegans* lives, deciding whether to leave or stay in a bacterial food patch when there is hydrogen peroxide in the environment is critical for survival.

Here, we show that hydrogen peroxide alters the bacterial food preference of *C. elegans*, enabling the nematodes to find food patches that provide hydrogen peroxide protection, where they can grow and reproduce. When H_2_O_2_ is present in the environment, the nematodes are more likely to leave food patches of laboratory and microbiome bacteria if those bacteria lack enzymes necessary for the degradation of environmental H_2_O_2_. This change in nematode behavior occurs because when bacterial communities have insufficient H_2_O_2_-degrading capacity, environmental H_2_O_2_ can excite the ASJ sensory neurons that promote escape from H_2_O_2_ and can prevent the response to bacteria of multiple classes of sensory neurons that promote locomotion towards bacteria. Thus, the modulation of *C. elegans’* sensory perception by the interplay of hydrogen peroxide and bacteria adjusts the nematode’s behavior to improve the nematode’s chances of finding a niche that provides both food and protection from hydrogen peroxide.

## Results

### Hydrogen peroxide alters the bacterial food preference of *C. elegans*

The bacterium *E. coli*, the food source of *C. elegans* under standard laboratory conditions, degrades environmental H_2_O_2_ primarily by expressing two catalases, KatG and KatE. These enzymes account for over 95% of *E. coli’s* H_2_O_2_-degrading capacity. The peroxiredoxin, AhpCF, plays a minor role (Seaver and Imlay, 2001). Previously, we found that *E. coli* JI377, a *katG katE ahpCF* triple null mutant strain which cannot degrade H_2_O_2_ in the environment (Seaver and Imlay, 2001), did not protect *C. elegans* adults from 1 mM H_2_O_2_ killing, whereas the *E. coli* MG1655 parental wild-type strain was protective (Schiffer et al., 2020). We observed a similar pattern when we quantified the development of *C. elegans* embryos in the presence or absence of 1 mM H_2_O_2_ in the environment: when no H_2_O_2_ was present, most embryos cultured on petri plates with either *E. coli* MG1655 or *E. coli* JI377 lawns developed into fertile adults (Figure S1). Embryos on plates with *E. coli* MG1655 and H_2_O_2_ also developed into fertile adults (Figure S1); however, those on plates with *E. coli* JI377 and H_2_O_2_ did not develop into adulthood and instead died as first stage (L1) larvae, similar to embryos that hatched on H_2_O_2_ plates without food (Figure S1). These findings showed that H_2_O_2_-degrading enzymes from *E. coli* created an environment where *C. elegans* was safe from the threat of H_2_O_2_, enabling the nematode’s development and subsequent reproduction.

Given the threat of H_2_O_2_ to *C. elegans* development and reproduction, we set out to determine the extent to which *C. elegans* would modulate their behavior to find an environment with both food and safety from the threat of H_2_O_2_. To determine whether *C. elegans* preferred *E. coli* strains that could degrade H_2_O_2_, we quantified the migration of populations of adult nematodes in a binary choice assay towards MG1655 and JI377 lawns (10^8^ bacterial cells each) on opposite sides of a petri plate (Figure 1A). In this assay, a choice index of 1 indicated complete preference for MG1655, a choice index of -1 indicated complete preference for JI377, and a choice index of 0 indicated no preference (Zhang et al., 2005). In the absence of added H_2_O_2_ in the environment, the nematodes moved toward both MG1655 and JI377 (Figure 1A). The fraction of nematodes on each lawn approximated a steady state after 30 minutes (Figure 1B). The nematodes exhibited a slight preference for MG1655: a choice index of 0.20 at the end of the two-hour assay (Figure 1C). In contrast, in the presence of 1 mM H_2_O_2_ in the environment, the nematodes showed a strong preference for MG1655 (a final choice index of 0.63, Figure 1C). We conclude that H_2_O_2_ altered the *E. coli* preference of *C. elegans*, increasing the nematode’s chances of finding an *E. coli* lawn that degrades H_2_O_2_.

**Figure 1.**
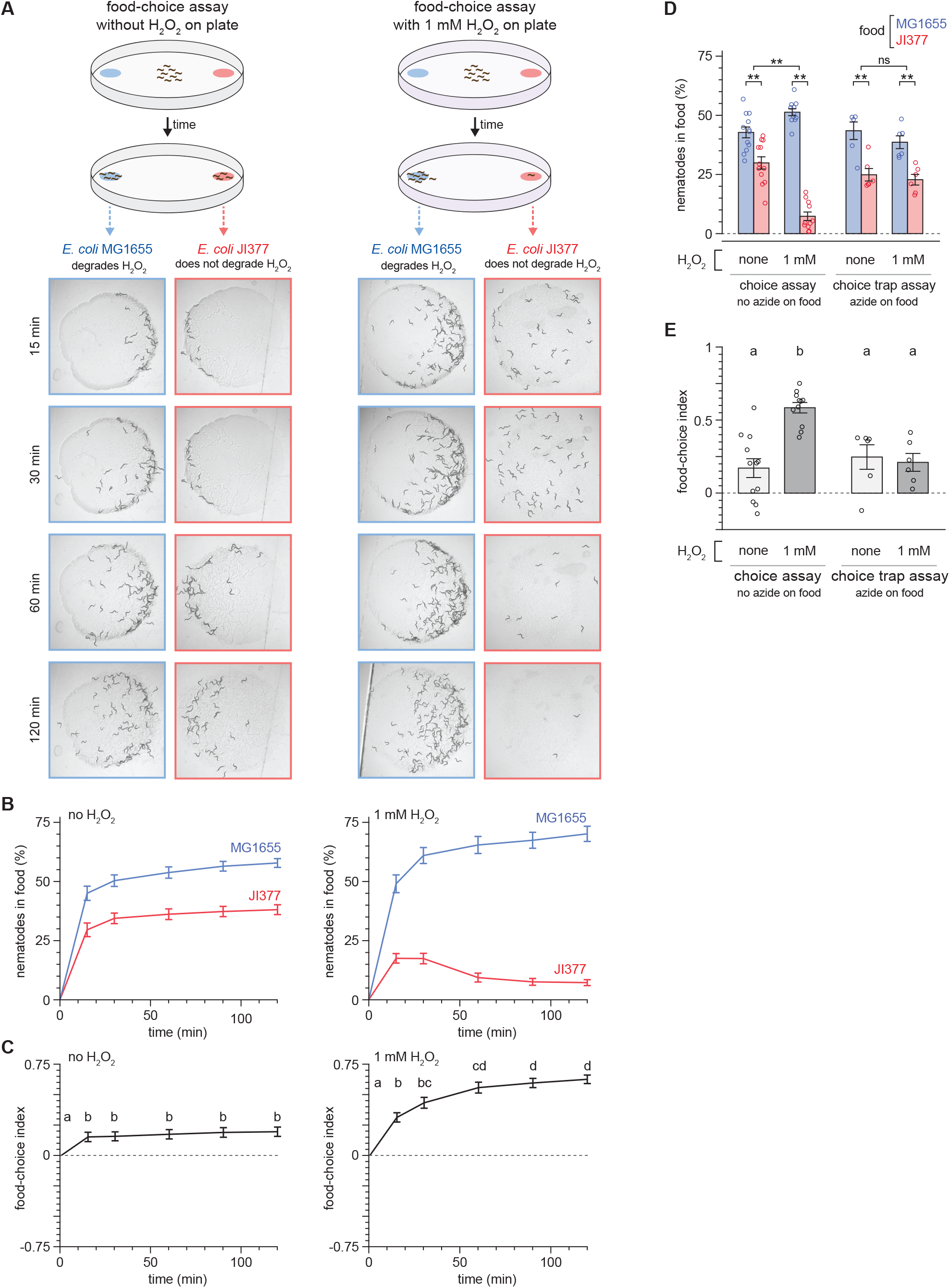
Hydrogen peroxide alters the bacterial food preference of *C. elegans*. (A) Diagram summarizing experimental strategy (top) and series of pictures of the *E. coli* MG1655 and JI377 lawns at the specified timepoints from representative food-choice assays (bottom) without added H_2_O_2_ (left) and with 1 mM H_2_O_2_ (right). (B) The proportion of nematodes on the *E. coli* MG1655 and JI377 lawns in assays without added H_2_O_2_ (left) and with 1 mM H_2_O_2_ (right) is plotted against time. *P* < 0.001 for times other than zero (ANOVA). n ≥ 15 assays per condition. (C) The food-choice indices for the assays shown in (B) are plotted against time. H_2_O_2_ induced an increase in food-choice index. Groups labeled with different letters exhibited significant differences (*P* < 0.05, Tukey HSD test) otherwise (*P* > 0.05). (D) The H_2_O_2_-dependent increase in the proportion of nematodes on the *E. coli* MG1655 lawns compared to JI377 lawns in two-hour food-choice assays was absent in choice-trap assays, in which the paralytic agent sodium azide was added to the bacterial lawns. ** indicates *P* < 0.002 and “ns” indicates *P* > 0.05 (standard least-squares regression). (E) The H_2_O_2_-induced increase in food-choice index was absent in choice-trap assaysn for the assays shown in (D). Groups labeled with different letters exhibited significant differences (*P* < 0.01, Tukey HSD test) otherwise (*P* > 0.05). Data are represented as mean ± s.e.m.

The increased nematode preference for *E. coli* MG1655 in the presence of H_2_O_2_ appeared to be in part due to an H_2_O_2_-induced change in nematode behavior after reaching the *E. coli* JI377 lawn. Instead of staying on the JI377 lawn as they did on assays without H_2_O_2_, in the presence of H_2_O_2_ a large proportion of nematodes left the JI377 lawn (Figure 1A); as a result, the fraction of nematodes on the JI377 lawn peaked 15 minutes after the start of the assay and decreased 2.4-fold thereafter, while the fraction of nematodes on the MG1655 lawn continued to increase (Figure 1B). To test whether the increased preference for MG1655 in the presence of H_2_O_2_ was due to an H_2_O_2_-dependent increase in the proportion of nematodes that left the JI377 lawn after reaching it, we used the paralytic agent sodium azide to prevent nematodes from leaving the bacterial lawns that they reached. Under these conditions, environmental H_2_O_2_ no longer increased the nematodes’ preference for MG1655 (Figure 1D-E). These findings suggested that environmental H_2_O_2_ increased *C. elegans* preference for *E. coli* lawns that degraded H_2_O_2_ primarily by increasing the chances that nematodes would leave lawns that did not degrade environmental H_2_O_2_.

### The H_2_O_2_-degrading capacity of bacterial communities determines nematode food leaving in response to environmental H_2_O_2_

To study how H_2_O_2_ induced *C. elegans* to leave an environment where food was plentiful, we used a bacterial lawn-leaving assay. In this assay, L4 stage nematodes were placed on a lawn with 10^8^ bacteria on a petri plate, and the proportion of nematodes leaving the lawn was measured after two hours (Figure 2A). Very few nematodes left either *E. coli* MG1655 or *E. coli* JI377 lawns when no H_2_O_2_ was added (Figure 2B-C). With 1 mM H_2_O_2_ in the environment, the proportion of nematodes leaving *E. coli* JI377 lawns increased relative to the control group (no added H_2_O_2_), but the proportion of nematodes leaving *E. coli* MG1655 lawns was unaffected (Figures 2B-C). We observed a similar H_2_O_2_-induced food leaving behavior in day 1 adults (Figure S2A). Pre-treating the *E. coli* JI377 suspension that we used to make the lawn with 1 mM H_2_O_2_ for 20 hours did not increase nematode lawn leaving when no H_2_O_2_ was added to the assay plates (Figure S2B). We conclude that the H_2_O_2_-induced food leaving behavior of *C. elegans* was caused by the failure of *E. coli* JI377 to degrade H_2_O_2_ in the environment.

**Figure 2.**
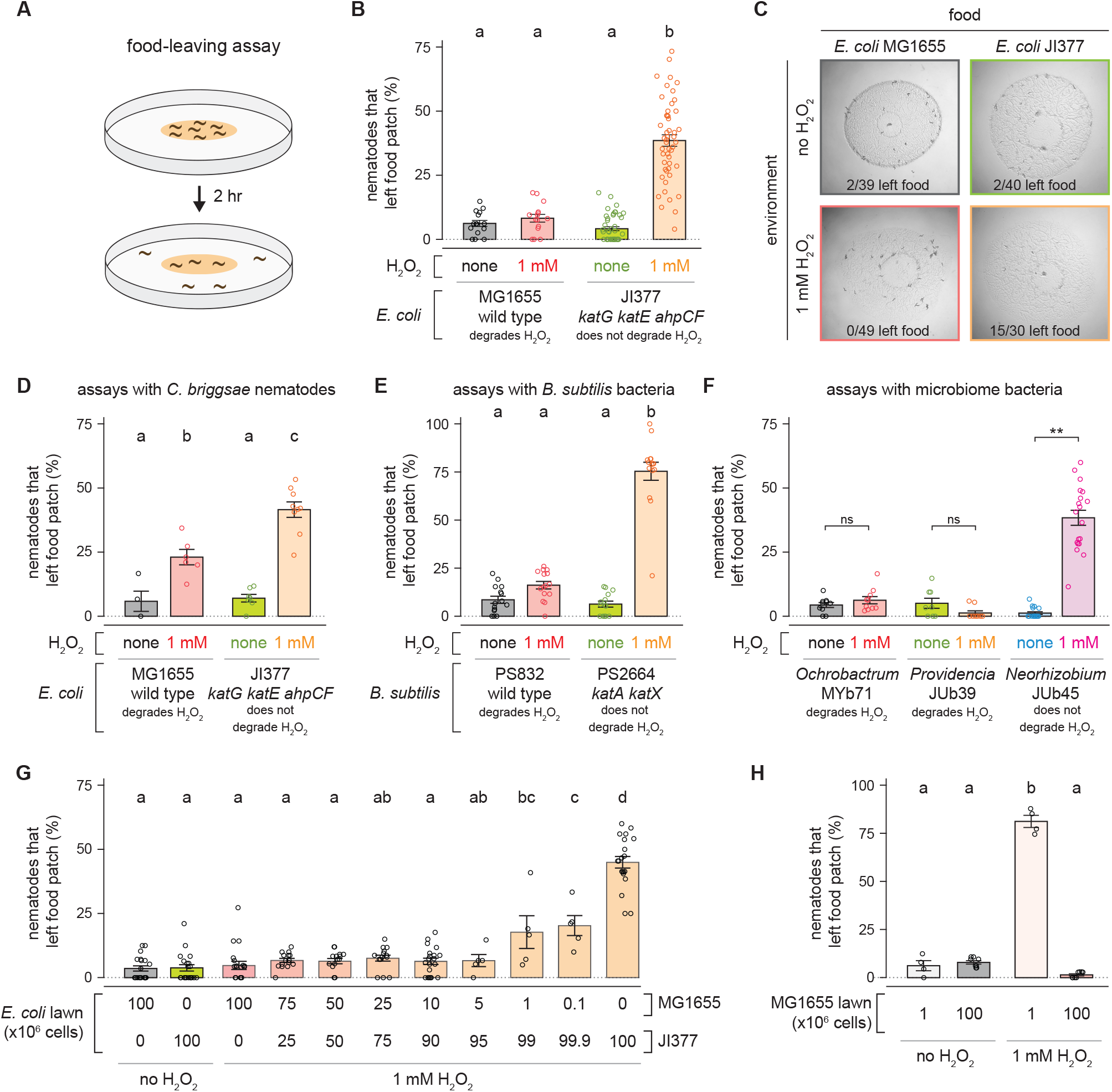
The H_2_O_2_-degrading capacity of bacterial communities determines nematode food leaving in response to environmental H_2_O_2_. (A) Diagram summarizing experimental strategy. (B) H_2_O_2_ induced an increase in the proportion of *C. elegans* nematodes that left a patch of *E. coli* JI377 but not of *E. coli* MG1655. (C) Representative pictures of the food-leaving assays shown in panel (B). (D) H_2_O_2_ induced a larger increase in the proportion of *C. briggsae* nematodes that left a patch of *E. coli* JI377 than of *E. coli* MG1655. (E) H_2_O_2_ induced an increase in the proportion of *C. elegans* nematodes that left a patch of *B. subtilis* PS2664 but not of *B. subtilis* PS832. (F) H_2_O_2_ induced an increase in the proportion of *C. elegans* nematodes that left a patch of *Neorhizobium* JUb45 but not of *Ochrobactrum* MYb71 or *Providencia* JUb39. ** indicates *P* < 0.0001 and “ns” indicates *P* > 0.05 (ANOVA). (G) H_2_O_2_-induced food leaving is determined by the total H_2_O_2_-degrading capacity of the *E. coli* patch. (H) H_2_O_2_-induced food leaving is determined by the number of *E. coli* MG1655 in the patch. Data are represented as mean ± s.e.m. Groups labeled with different letters exhibited significant differences (*P* < 0.05, Tukey HSD test) otherwise (*P* > 0.05).

To determine whether the H_2_O_2_-induced food leaving behavior we observed in *C. elegans* was conserved in other nematodes, we repeated the food-leaving assays using *Caenorhabditis briggsae*, a nematode species that diverged from *C. elegans* approximately 100 million years ago (Hillier et al., 2007). We found that the proportion of *C. briggsae* nematodes that left MG1655 lawns was higher on plates with 1 mM H_2_O_2_ (Figure 2D). In assays with JI337, H_2_O_2_ induced a larger increase in in the proportion of *C. briggsae* nematodes that left the lawn than in assays with MG1655 (Figure 2D). Therefore, H_2_O_2_ induced food leaving in two distantly related nematode species.

Next, we determined whether the induction of nematode food leaving by H_2_O_2_ was specific to *E. coli*, a Gram-negative bacterium, or extended to other bacteria. The Gram-positive bacterium *Bacillus subtilis* has multiple catalase genes: *katA*, expressed in vegetative cells (Bol and Yasbin, 1994), and *katX*, expressed in spores (Casillas-Martinez and Setlow, 1997). We found that when no H_2_O_2_ was added, fewer than 10% of the nematodes left wild-type or *katA katX* double mutant *B. subtilis* lawns (Figure 2E). With 1 mM H_2_O_2_ in the environment, the proportion of nematodes leaving *katA katX B. subtilis* lawns increased to 75% but the proportion of nematodes leaving wild-type *B. subtilis* lawns was not affected (Figure 2E). Therefore, the H_2_O_2_-degrading capacity of both Gram-positive and Gram-negative bacteria determined nematode food leaving in response to environmental H_2_O_2_.

*C. elegans* encounters a wide variety of bacterial taxa in its natural habitat (Samuel et al., 2016; Zhang et al., 2017). We first determined whether genes encoding H_2_O_2_-degrading enzymes were present in the sequenced genomes of 180 strains isolated from *C. elegans* habitats (Table S1). While all strains possessed at least one gene encoding an H_2_O_2_-degrading enzyme, they exhibited a wide range in the number (from 3 in *Lactococcus lactis* BIGb0220 to 31 in *Sphingobacterium* JUb78) and types of H_2_O_2_-degrading enzymes they encoded (Figure S3A-B), suggesting potential variation in H_2_O_2_-degrading capacities of these strains.

To directly assess the extent to which individual bacterial species from natural habitats of *C. elegans* can degrade environmental H_2_O_2_, we measured the catalase activity of 165 strains isolated from *C. elegans* habitats, including 102 strains from compost microcosms, 39 from various rotting fruits, and 23 from snails and slugs (Table S2). Most of the strains we screened exhibited catalase activity, including the 12 strains from 9 families in the CeMbio “core microbiome” collection (Dirksen et al., 2020). However, seventeen strains lacked or had very poor catalase activity (Table S2), including *Neorhizobium sp*. JUb45 (from a rotting apple), consistent with a previous study (Kniazeva and Ruvkun, 2019), *Lactococcus lactis* BIGb0220 (from a rotting apple), consistent with the lack of catalase genes in its genome (Table S1), one of six *Microbacterium* strains (JUb76, from a rotting grape), and all 14 *Shewanella* strains (from compost and defining at least four independent isolates based on 16S rRNA gene sequence, Figure S4). Catalases are commonly induced as bacterial cells enter stationary phase, because these enzymes degrade hydrogen peroxide without depleting cellular energy sources (Mishra and Imlay, 2012). *B. subtilis* PS832 induced catalase expression during the transition from exponential growth to stationary phase (Table S2), consistent with previous work (Bol and Yasbin, 1994). Similarly, *Microbacterium* JUb76 showed growth phase-dependent catalase activity: the strain lacked catalase activity in exponential growth phase but had low catalase activity during stationary phase (Table S2). We next sought to determine whether sequence variation in the catalase KatE and KatG orthologs could explain changes in observed functions. The *Neorhizobium* sp. JUb45 genome encodes one *katE* and one *katG* ortholog (Table S2). Alignments of the predicted proteins to their *E. coli* orthologs indicated general conservation of the predicted catalytic residues for both KatE [His128, Asn201, and Tyr415 (Loewen, 1996)] and KatG [Arg102, Trp105, His106, and His267 (Powers et al., 2001)] (Figure S3C). However, H_2_O_2_ ligand binding domain residues exhibited much more variation between *E. coli* and several microbiome members including *Neorhizobium sp*. JUb45 (Figure S3C). All together, these studies showed that both catalase positive and negative bacteria were common in natural habitats of *C. elegans* and suggested these nematodes may encounter bacteria that exhibit a poor H_2_O_2_-degrading capacity.

We next determined whether the H_2_O_2_-degrading capacity of beneficial bacteria isolated from natural habitats of *C. elegans* predicted the food-leaving behavior of *C. elegans* in the presence of environmental H_2_O_2_. Most nematodes did not leave lawns of catalase-positive *Ochrobactrum vermis* MYb71 (from the CeMbio collection) and *Providencia alcalifaciens* JUb39 even when H_2_O_2_ was added (Figure 2F). In contrast, with H_2_O_2_ in the environment, about half of the nematodes left lawns of catalase-negative *Neorhizobium sp*. JUb45, a much higher proportion than when no H_2_O_2_ was added (Figure 2F). These studies suggested that the H_2_O_2_-degrading capacity of microbiome species may determine the food leaving behavior of nematodes in their natural habitat.

In the natural environment *C. elegans* is unlikely to encounter homogeneous bacterial lawns consisting of a single bacterial genotype. To investigate how the H_2_O_2_-degrading capacity of a lawn influences the nematode’s H_2_O_2_-induced food-leaving behavior, we varied the composition of the bacterial lawns by mixing varying proportions of *E. coli* MG1655 and JI377 while keeping constant the total number of bacteria within the lawn (10^8^ cells). We found that when 1 mM H_2_O_2_ was added, the nematodes did not leave lawns with 0.1% MG1655 and 99.9% JI377 as much as they left lawns with only JI377 (Figure 2G). Therefore, the nematode’s food-leaving behavior was highly sensitive to the proportion of *E. coli* cells in a lawn that were able to degrade H_2_O_2_ in the environment. The total number of *E. coli* cells in the lawn was also important, because when H_2_O_2_ was added most nematodes left a MG1655 lawn with just 10^6^ cells (Figure 2H), even though that number of MG1655 cells was sufficient to retain the nematodes in a lawn with a total of 10^8^ cells composed of 1% MG1655 and 99% JI377 (Figure 2G). Taken together, these findings suggested that the nematode’s H_2_O_2_-induced food leaving is determined by the bacterial community’s population size and total H_2_O_2_-degrading capacity.

### Production of serotonin inhibits H_2_O_2_-induced food leaving

Because food levels affected the H_2_O_2_-induced food leaving behavior of *C. elegans*, we speculated that this behavior may be regulated by the neurotransmitter serotonin. Expression of the serotonin biosynthetic tryptophan hydroxylase gene *tph-1* increases with food (Cunningham et al., 2012; Entchev et al., 2015; Liang et al., 2006; Patel et al., 2020; Sze et al., 2000; Zheng et al., 2005) and serotonin regulates many food-related behaviors (Liang et al., 2006; Melo and Ruvkun, 2012; Sze et al., 2000; Zheng et al., 2005). We found that *tph-1(mg280)* null mutants, which specifically lack serotonin (Sze et al., 2000), were more likely than wild-type animals to leave a lawn of *E. coli* JI377 when 1 mM H_2_O_2_ was added to the environment (Figure 3A). The *tph-1(mg280)* mutation did not affect either the proportion of nematodes leaving a lawn of *E. coli* JI377 when no H_2_O_2_ was added, or the proportion of nematodes leaving a lawn of *E. coli* MG1655 even when H_2_O_2_ was added (Figure 3A). A second, independently derived, *tph-1(n4622)* null deletion allele (Shivers et al., 2009), exhibited the same behavior as *tph-1(mg280)* (Figure 3B). Therefore, serotonin biosynthesis inhibits *C. elegans’* decision to leave a food lawn that does not provide H_2_O_2_ protection.

**Figure 3.**
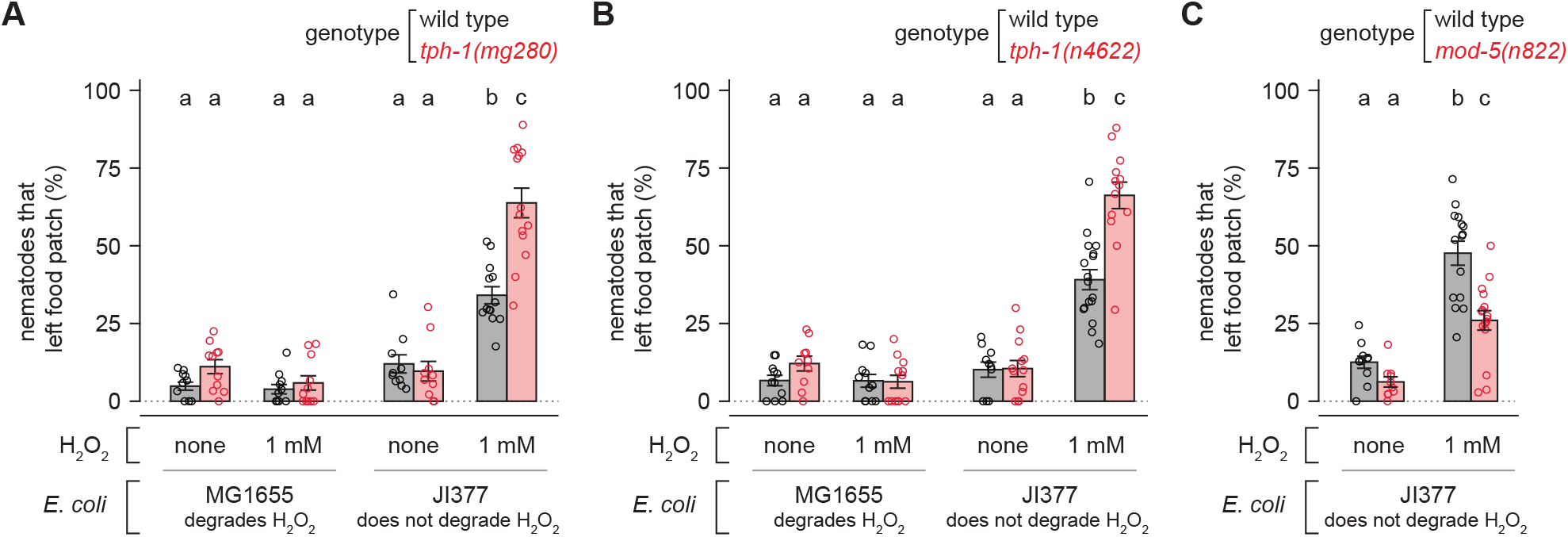
Production of serotonin inhibits H_2_O_2_-induced food leaving. (A-B) *tph-1* null mutations increase the proportion of nematodes that leave an *E. coli* JI377 food patch in the presence of H_2_O_2_. (C) A *mod-5* null mutation decreases the proportion of nematodes that leave an *E. coli* JI377 food patch in the presence of H_2_O_2_. Data are represented as mean ± s.e.m. Groups labeled with different letters exhibited significant differences (*P* < 0.05, Tukey HSD test) otherwise (*P* > 0.05).

Since lowering serotonin levels increased food-leaving when the *E. coli* lawn did not degrade environmental H_2_O_2_, we determined whether increasing serotonin levels would be sufficient to lower food leaving. Nematodes with a serotonin reuptake transporter gene *mod-5(n822)* null mutation have higher presynaptic serotonin levels (Ranganathan et al., 2001). We found that *mod-5(n822)* mutants were less likely to leave an *E. coli* JI377 lawn than wild-type animals (Figure 3C). We conclude that serotonin functions in a dose-dependent manner to inhibit the nematode’s H_2_O_2_-induced food leaving behavior.

### Hydrogen peroxide and bacteria have opposing effects on the activity of sensory neurons

How did *C. elegans* overcome its strong attraction towards *E. coli* to specifically leave lawns that did not degrade hydrogen peroxide? *C. elegans* relies on sensory perception of bacterially derived cues to efficiently find the bacteria it feeds on (Ferkey et al., 2021). Most sensory functions in *C. elegans* hermaphrodites are performed by 60 ciliated and 12 non-ciliated neurons (White et al., 1986). Twelve pairs of those ciliated neurons make up the nematode’s major sensory organs, the two amphids, responsible for smell, taste, and temperature sensation (Bargmann, 2006). To assess the extent to which *E. coli* and H_2_O_2_ modulated the function of amphid sensory neurons, we examined their activity in response to combinations of *E. coli* and H_2_O_2_.

The activity of *C. elegans* sensory neurons is strongly correlated with their calcium responses (Ramot et al., 2008). We presented nematodes expressing the genetically encoded calcium indicator GCaMP6 in sensory neurons with six combinations of stimuli consisting of suspensions of *E. coli* MG1655, *E. coli* JI377, or water, pre-mixed with or without 1 mM H_2_O_2_ for 20 hours (to give the bacteria an opportunity to break down the H_2_O_2_). We used a custom-built microfluidic device to deliver these stimuli to the amphids of each L4 stage nematode in a randomized order, in 15 second intervals, preceded and followed by 45 second intervals without stimuli, while recording with single-cell resolution the activity of 26 sensory neurons via fluorescence microscopy (Figure 4A-D). Our imaging studies covered 11 of the 12 pairs of amphid neurons (ADF, ADL, ASE, ASG, ASH, ASI, ASJ, ASK, AWA, AWB, and AWC), and 2 pairs of non-amphid neurons (BAG and URX). The six combinations of stimuli induced two major patterns of neuronal modulation, described below, with each pattern affecting the activity of multiple sensory neurons: one pattern was induced by 1 mM H_2_O_2_ and *E. coli* JI377 with 1 mM H_2_O_2_, and the other pattern was induced by *E. coli* MG1655, *E. coli* JI377, and *E. coli* MG1655 with 1 mM H_2_O_2_ (Figure 4E).

**Figure 4.**
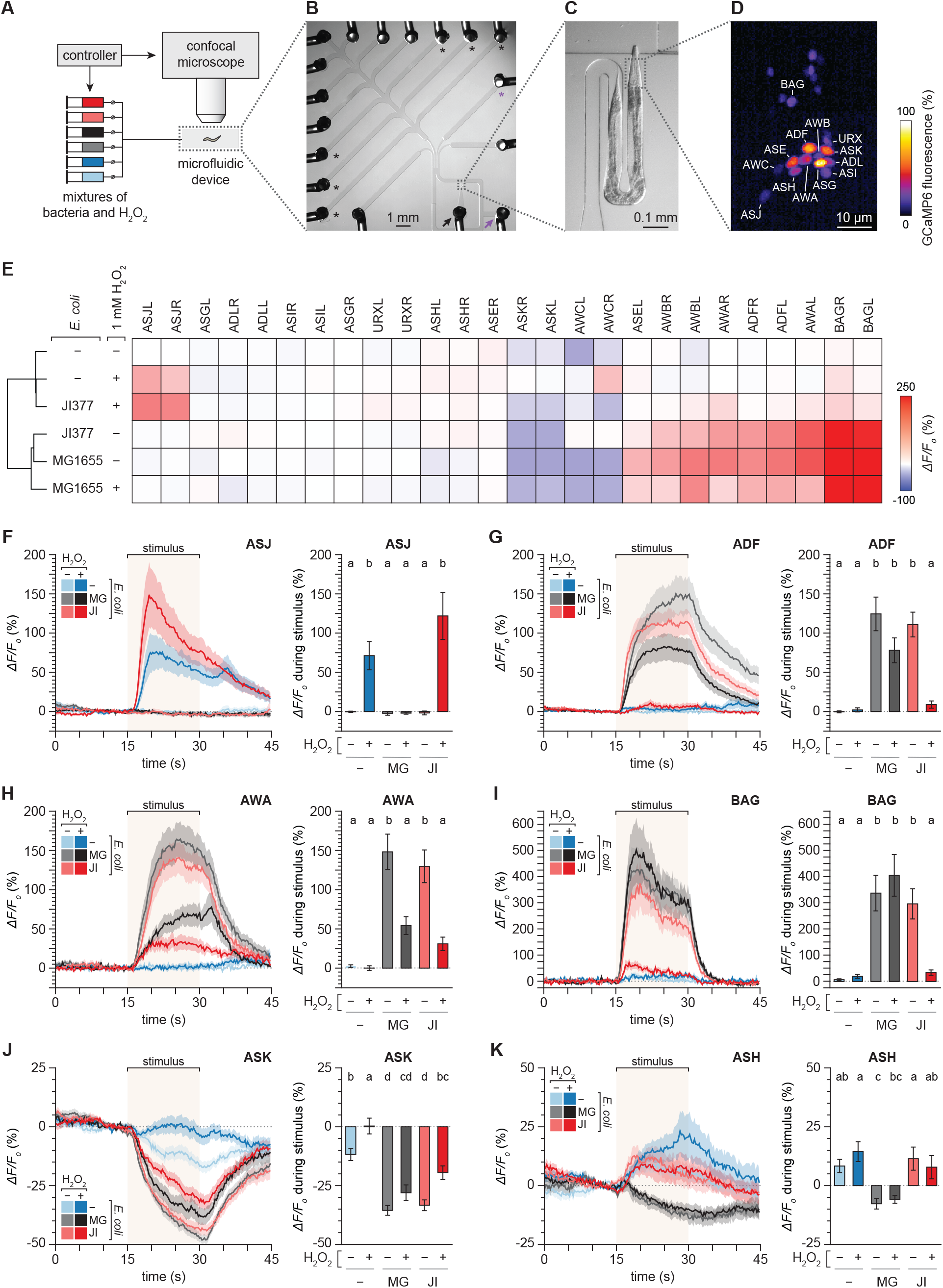
Hydrogen peroxide and bacteria have opposing effects on the activity of sensory neurons. (A) Schematic of the microfluidic setup for controlled delivery of sensory stimuli and calcium imaging of nematodes. (B) Multi-channel microfluidic device. The black arrow marks the inlet channel for loading the nematode, the purple arrow marks the outlet channel for fluid waste, the black asterisks mark the stimuli delivery channels that were used, and the purple asterisk marks the buffer delivery channel. (C) Magnified view of the channel where the sensory endings in the head of the immobilized nematode are stimulated with mixtures of bacteria and H_2_O_2_. (D) Relative fluorescence of the GCaMP6 genetically encoded calcium indicator expressed by the animal. 26 sensory neurons in the nematode’s head (13 shown) were imaged with single cell resolution. (E) Clustering of the mean changes in GCaMP6 fluorescence of 26 sensory neurons in response to six stimuli consisting of suspensions of *E. coli* MG1655, *E. coli* JI377, or water, with or without adding 1 mM H_2_O_2_. (F-K) Average GCaMP6 fluorescence traces of (F) ASJ, (G) ADF, (H) AWA, (I) BAG, (J) ASK, and (K) ASH neuronal classes in response to six different stimuli (left sub-panels) and average changes in fluorescence in response to those stimuli (right sub-panels). The stimulus delivery interval is indicated by a shaded box. Data are represented as mean ± s.e.m. The number of neurons imaged was 28 ADF, 28 ADL, 14 ASEL, 14 ASER, 28 ASG, 28 ASH, 27 ASI, 28 ASJ, 28 ASK, 28 AWA, 13 AWB, 24 AWC, 18 BAG, and 28 URX. Groups labeled with different letters exhibited significant differences (*P* < 0.05, Tukey HSD test) otherwise (*P* > 0.05). Traces for the URX, ADL, ASEL, ASER, ASG, ASI, AWB, and AWC neuronal classes are shown in Figure S4.

H_2_O_2_ strongly excited (increased [Ca^2+^]) the ASJ neuron pair (Figure 4F) which, later in this manuscript, we show is required for H_2_O_2_ avoidance. In the ASK and URX neuron pairs, H_2_O_2_ also increased the GCaMP6 signal relative to the water-only control, albeit more weakly than in the ASJ neuron pair (Figures 4J and S5A). These H_2_O_2_-induced increases in neuronal activity were abolished when H_2_O_2_ was combined with *E. coli* MG1655 (which degrades H_2_O_2_) but persisted when H_2_O_2_ was combined with *E. coli* JI377 (which does not degrade H_2_O_2_) (Figures 4F,J and S5A). Therefore, *E. coli’s* H_2_O_2_-degrading enzymes prevented the excitation of ASJ, ASK, and URX neurons by environmental H_2_O_2_.

*E. coli* MG1655 and JI377 modulated the activity of mostly overlapping but distinct sets of sensory neurons. Both *E. coli* strains excited the ADF, AWA, AWB, and BAG neuron pairs (Figures 4G-I and S5G) and inhibited (decreased [Ca^2+^]) the ASK pair (Figure 4J). MG1655 also excited ASEL (Figure S5C) and inhibited the ASH pair (Figure 4K), while JI377 excited the ADL pair (Figure S5B). When combined with H_2_O_2_, MG1655 elicited the same response pattern in ADF, ASK, ASH, and BAG as it did without H_2_O_2_ (Figure 4G,I-K). In contrast, when combined with H_2_O_2_, JI377 no longer elicited a significant response in ADF, ADL, ASK, AWA, and BAG (Figures 4G-K and S5B). Therefore, *E. coli’s* H_2_O_2_-degrading enzymes prevented H_2_O_2_ from blocking the specific excitation or inhibition of most of the sensory neurons that were modulated by *E. coli*.

Most of the neuronal classes excited by *E. coli* MG1655 mediate locomotion towards attractive cues. ADF and ASEL sense water-soluble attractants (Bargmann and Horvitz, 1991; Pierce-Shimomura et al., 2001), AWA detect attractive volatile odorants (Bargmann et al., 1993), and BAG sense oxygen and carbon dioxide (Guillermin et al., 2017; Zimmer et al., 2009). The neuronal classes inhibited by *E. coli* MG1655 mediate locomotion away from repulsive cues; ASK and ASH sense various repellents (Gray et al., 2005; Hilliard et al., 2002). We propose that, when *E. coli* cannot degrade environmental H_2_O_2_, the strong attraction of *C. elegans* to *E. coli* is weakened because H_2_O_2_ prevents the modulation of those classes of neurons by *E. coli*, increasing the chances that the nematode would leave the *E. coli* lawn.

### The H_2_O_2_-sensing ASJ neurons are required for H_2_O_2_ avoidance

Because the ASJ neuronal pair was strongly excited by H_2_O_2_ (Figure 4F), we speculated these neurons may mediate an aversive locomotory response to H_2_O_2_, in line with the role of ASJ in triggering an aversive response when excited by cues from the *C. elegans* predator *Pristionchus pacificus* (Liu et al., 2018) and from the bacterial pathogen *Pseudomonas aeruginosa* (Meisel et al., 2014). To quantify H_2_O_2_ avoidance, we exposed nematodes to a drop of 1 mM H_2_O_2_ or water and recorded the proportion of avoidance responses (a reversal followed by an omega bend, Figure 5A) in response to these stimuli (Broekmans et al., 2016; Hilliard et al., 2002; Liu et al., 2018). H_2_O_2_ elicited a significant increase in the proportion of avoidance responses relative to the water control (Figure 5B). In animals in which the ASJ neurons were genetically ablated via ASJ-specific caspase expression (Cornils et al., 2011), that increase was absent (Figure 5C). Therefore, the H_2_O_2_-sensing ASJ neurons were required for H_2_O_2_ avoidance. This ASJ-dependent aversive response to H_2_O_2_ enables *C. elegans* to escape environments with lethal H_2_O_2_ levels.

**Figure 5.**
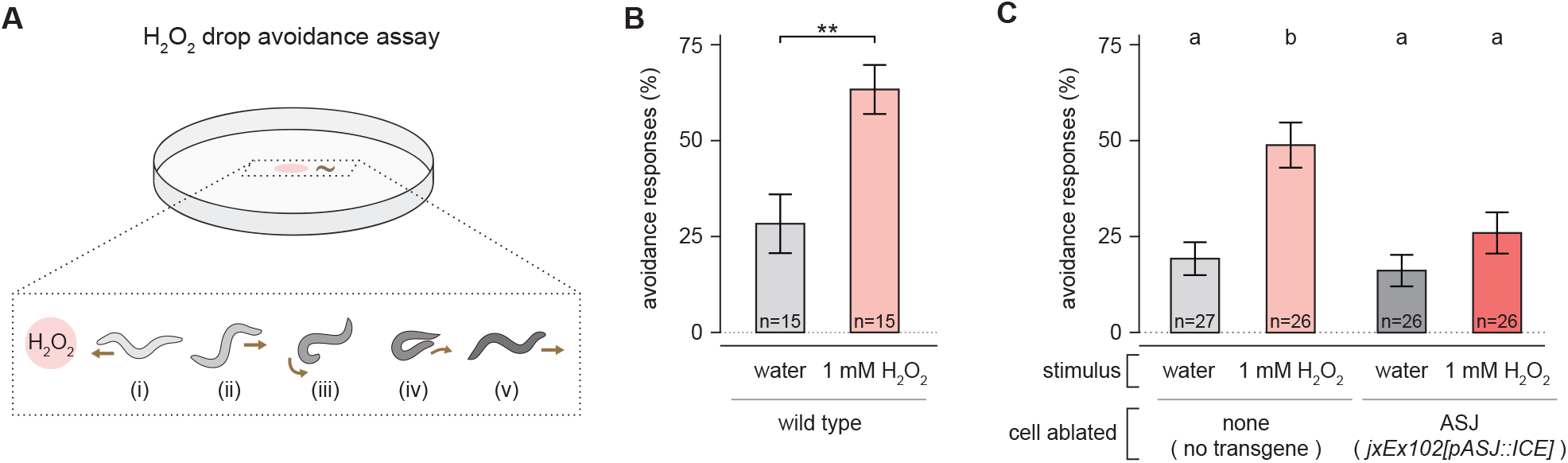
The H_2_O_2_-sensing ASJ neurons are required for H_2_O_2_ avoidance. (A) Schematic overview of hydrogen peroxide drop avoidance assay, with worm body shapes extracted from tracking data (Donnelly et al., 2013): upon sensing the small volume of 1 mM H_2_O_2_ on its path (i), the nematode initiates an avoidance response consisting of a reversal phase (ii), an omega turn (iii-iv), and the resumption of locomotion (v). (B) H_2_O_2_ induces an increase in avoidance responses. ** indicates *P* < 0.002 (t-test). (C) Ablation of the ASJ neurons suppresses the increase in avoidance responses induced by H_2_O_2_. Groups labeled with different letters exhibited significant differences (*P* < 0.01, Tukey HSD test) otherwise (*P* > 0.05). Data are represented as mean ± s.e.m. of the average avoidance response of each animal per condition.

### Hedging the H_2_O_2_-induced food leaving decision provides an adaptation to changing environments

We reasoned that *C. elegans’* decision to leave a bacterial lawn that did not degrade lethal concentrations of H_2_O_2_ was adaptive, because it would have given the nematodes a chance to find a safe environment with food conducive to their reproduction and the survival of their progeny. We were, therefore, surprised to find that a lethal concentration of H_2_O_2_ did not induce all nematodes in the population to leave an *E. coli* lawn unable to degrade H_2_O_2_; instead, a large proportion of nematodes remained in the lawn even after two hours (Figure 2B). Because the evolution of adaptive behaviors is thought to be shaped by the stability of the organism’s environment (Gordon, 2016), we reasoned that perhaps remaining on the *E. coli* lawn would be adaptive when adverse conditions are temporary.

To explore that possibility, we examined the nematode’s H_2_O_2_-induced food-leaving behavior over a longer timescale. When no H_2_O_2_ was added, few nematodes left the *E. coli* JI377 lawn over a four-hour period. With 1 mM H_2_O_2_ in the environment, the proportion of nematodes that left the *E. coli* JI377 lawn reached a steady state after increasing for the first two hours (Figure 6A). To investigate why the nematodes remained on the *E. coli* lawn after two hours even though H_2_O_2_ was present, we examined their locomotory behavior throughout the food-leaving assay. When no H_2_O_2_ was added, most nematodes were roaming (moving forward rapidly and turning infrequently) and a small proportion of the nematodes were dwelling (turning frequently or staying in place while feeding) (Figure 6B). In contrast, in the presence of 1 mM H_2_O_2_, the proportion of nematodes roaming steadily decreased and the proportion of nematodes in a quiescent state increased until most nematodes were quiescent after four hours (Figure 6B). Quiescence is a state of complete immobility and cessation of feeding (You et al., 2008) often induced by stressful environmental conditions (Hill et al., 2014). After two hours in the presence of H_2_O_2_ more than half the nematodes were roaming and 20% were dwelling (Figure 6B), and those proportions were the same whether the nematodes left or stayed in the bacterial lawn (Figure S6). Thus, after a two-hour exposure to H_2_O_2_ most of the nematodes that remained in the bacterial lawn were capable of leaving that lethal environment.

**Figure 6.**
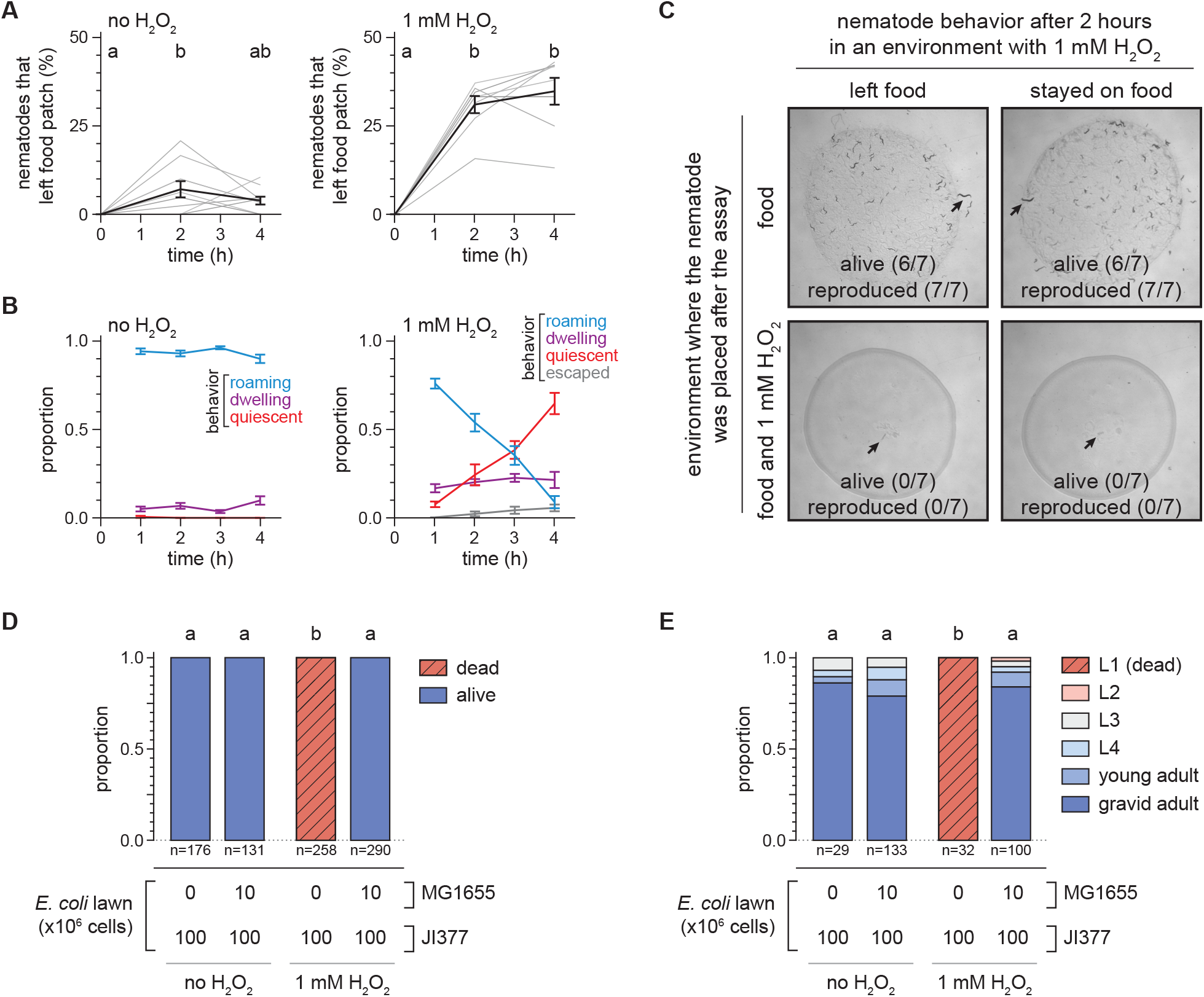
Hedging the H_2_O_2_-induced food leaving decision provides an adaptation to changing environments. (A) H_2_O_2_ induced an increase in the proportion of *C. elegans* nematodes that left a patch of *E. coli* JI377. Groups labeled with different letters exhibited significant differences (*P* < 0.05, Tukey HSD test) otherwise (*P* > 0.05). Data are represented as mean ± s.e.m of n ≥ 8 assays per condition (grey lines). (B) H_2_O_2_ induced time-dependent changes in the proportion of nematodes roaming, quiescent, or that escaped the petri plate, and induced increased nematode dwelling, in an *E. coli* JI377 food-leaving assay (*P* < 0.01, standard least-squares regression). Data are represented as mean ± s.e.m of n = 6 assays per condition. (C) Representative pictures of the survival and reproduction of individual wild-type *C. elegans* 72 hours after being transferred to plates with *E. coli* JI377 +/- 1 mM H_2_O_2_ from a food-leaving assay on JI377 with 1 mM H_2_O_2_. (D) Survival of wild-type *C. elegans* 72 hours after the end of a food-leaving assay. The *E. coli* MG1655 was added onto the lawn at the end of the food-leaving assay. Groups labeled with different letters exhibited significant differences (*P* < 0.0001, Fisher’s exact test) otherwise (*P* > 0.05). (E) Development of wild-type *C. elegans* embryos in the presence of 1 mM H_2_O_2_. The *E. coli* MG1655 was added onto the lawn immediately after the eggs were added. Groups labeled with different letters exhibited significant differences (*P* < 0.001, ordinal logistic regression) otherwise (*P* > 0.05).

Staying in a bacterial lawn in the presence of lethal levels of H_2_O_2_ may enable nematodes to wait for environmental conditions to become favorable for reproduction or could be a manifestation of irreversible damage to the nematodes caused by lethal levels of environmental H_2_O_2_. To distinguish between these possibilities, we transferred nematodes that had been in food-leaving assays for two hours to new environments. We found that if the nematodes were transferred from plates with JI377 and 1 mM H_2_O_2_ to new plates with JI377 but without H_2_O_2_, most nematodes survived and reproduced, whether they had left or stayed on the *E. coli* JI377 lawn (Figure 6C). In contrast, all nematodes transferred to new plates also with JI377 and H_2_O_2_ died after three days and had no progeny that reached adulthood (Figure 6C). We also found that nematode survival was restored if, at the end of a food leaving assay with 1 mM H_2_O_2_, we added 10^7^ MG1655 cells onto the JI377 lawn of 10^8^ cells (Figure 6D). We observed a similar pattern when we quantified the development of *C. elegans* embryos in the presence of 1 mM H_2_O_2_: the embryos died as L1 larvae after 72 hours on JI377 lawns of 10^8^ cells, but developed into fertile adults if after placing the eggs onto the JI377 lawn we added 10^7^ MG1655 cells to the lawn (Figure 6E). Because a small proportion of bacteria capable of degrading H_2_O_2_ could remodel an environment that was not conducive to the nematode’s survival and reproduction, we concluded that remaining in a bacterial lawn that contained a lethal concentration of environmental H_2_O_2_ was a plausible behavioral strategy for the nematode’s reproduction and the survival of their progeny. Based on these findings, we propose that the incomplete penetrance of *C. elegans* H_2_O_2_-induced food leaving could be understood as a bet-hedging adaptation to changing environments, because nematode reproduction occurred only if by leaving they found both food and safety or if when staying the environment turned favorable on its own.

## Discussion

In the present study, we developed a model ecosystem to study the behavioral mechanisms that enable the nematode *C. elegans* to find a niche that provides the food and safety necessary for growth and reproduction. We found that modulation of the nematode’s sensory perception by hydrogen peroxide—the most common chemical threat in the microbial battlefield (Mishra and Imlay, 2012)—enables the nematode to override its strong attraction towards the bacteria it feeds on and, thus, leave environments where the bacterial community does not provide the nematode and its future progeny with sufficient protection from hydrogen peroxide.

### *C. elegans* adjusts its behavior to find bacterial communities that provide protection from hydrogen peroxide

We show here that *C. elegans* adjusts its locomotory behavior in response to environmental H_2_O_2_; the nematode left niches where the bacterial community did not provide H_2_O_2_ protection and stayed in those that were protective. The induction of food leaving by H_2_O_2_ was determined by the H_2_O_2_-degrading capacity of Gram-negative *E. coli* and Gram-positive *B. subtilis*, both laboratory bacteria. The H_2_O_2_-induced food leaving behavior also occurred in *C. briggsae* nematodes, which are distantly related to *C. elegans*. These two nematode species feed on a wide variety of bacteria in their natural habitats (Avery and Shtonda, 2003; Felix and Duveau, 2012; Samuel et al., 2016). We found that while bacteria capable of degrading environmental H_2_O_2_ were common in natural habitats of *C. elegans*, these nematodes may often encounter bacteria with a poor H_2_O_2_-degrading capacity, including *Neorhizobium, Microbacterium*, and *Shewanella* species. The H_2_O_2_-degrading capacity of microbiome bacteria determined whether H_2_O_2_ induced *C. elegans* to leave food. In addition, the H_2_O_2_-induced food leaving behavior of *C. elegans* responded to the size and total H_2_O_2_-degrading capacity of the bacterial community. We propose that in their natural habitat, nematodes decide whether to feed on bacterial communities that provide sufficient food and protection from H_2_O_2_ or leave when they deem those communities do not provide sufficient food or H_2_O_2_ protection.

*C. elegans’* decision to leave a bacterial community that does not provide sufficient H_2_O_2_ protection is dictated by sensory neurons that respond to the perception of food and H_2_O_2_ in the environment (discussed in the next section). However, the outcome of this decision was not based solely on the perceived environmental conditions, as more than half the nematodes stayed on food lawns that did not provide protection from lethal H_2_O_2_ levels in the environment. The lack of unanimity in the choice of staying or leaving could be understood as a bet-hedging adaptation to changing environments, because leaving a lethal environment with food does not guarantee finding one conducive to growth and reproduction, while staying in an adverse environment may lead to survival and reproduction if conditions improve. Consistent with that possibility, nematodes that stayed on the bacterial lawns with lethal H_2_O_2_ levels survived and reproduced when environmental conditions improved with the addition of a small proportion of bacteria capable of degrading H_2_O_2_. We propose that the decision of not always leaving a food-rich but lethal environment provides the nematode with an evolutionarily optimal adaptive strategy (Kussell and Leibler, 2005; Maynard Smith, 1982; Wolf et al., 2005) to deal with the possibility that future environmental conditions may be more conducive to nematode growth and reproduction than the adverse conditions perceived by the nematode’s sensory neurons.

The decision to leave a bacterial lawn that does not provide H_2_O_2_ protection was also inhibited in a dose-dependent manner by the extracellular levels of serotonin, a neurotransmitter whose production and release is regulated in response to the sensory perception of food (Entchev et al., 2015; Iwanir et al., 2016; Rhoades et al., 2019; Song et al., 2013). Serotonin release slows nematode locomotion as they approach and encounter bacterial lawns (Iwanir et al., 2016; Rhoades et al., 2019). Therefore, serotonin may couple the nematode’s H_2_O_2_-induced food-leaving behavior to the level and quality of food in the environment.

### *C. elegans* assess faithfully the threat of hydrogen peroxide via sensory perception

How does *C. elegans* decide to leave an environment in which the bacterial community does not provide H_2_O_2_ protection? We propose that *C. elegans* accomplishes that task though the action of both H_2_O_2_-sensing neurons that promote escape from H_2_O_2_ and bacteria-sensing neurons that promote attraction to bacteria. The action of those H_2_O_2_-sensing neurons is modulated by *E. coli* H_2_O_2_-degrading enzymes, which degrade the input to those neurons. The action of those bacteria-sensing neurons is modulated by H_2_O_2_, which prevents their specific excitation or inhibition by *E. coli*. Together these sets of sensory neurons ensure the nematode’s behavior is faithful to the threat of H_2_O_2_, which is contingent on the bacterial capacity to degrade H_2_O_2_ (Figure 7).

**Figure 7.**
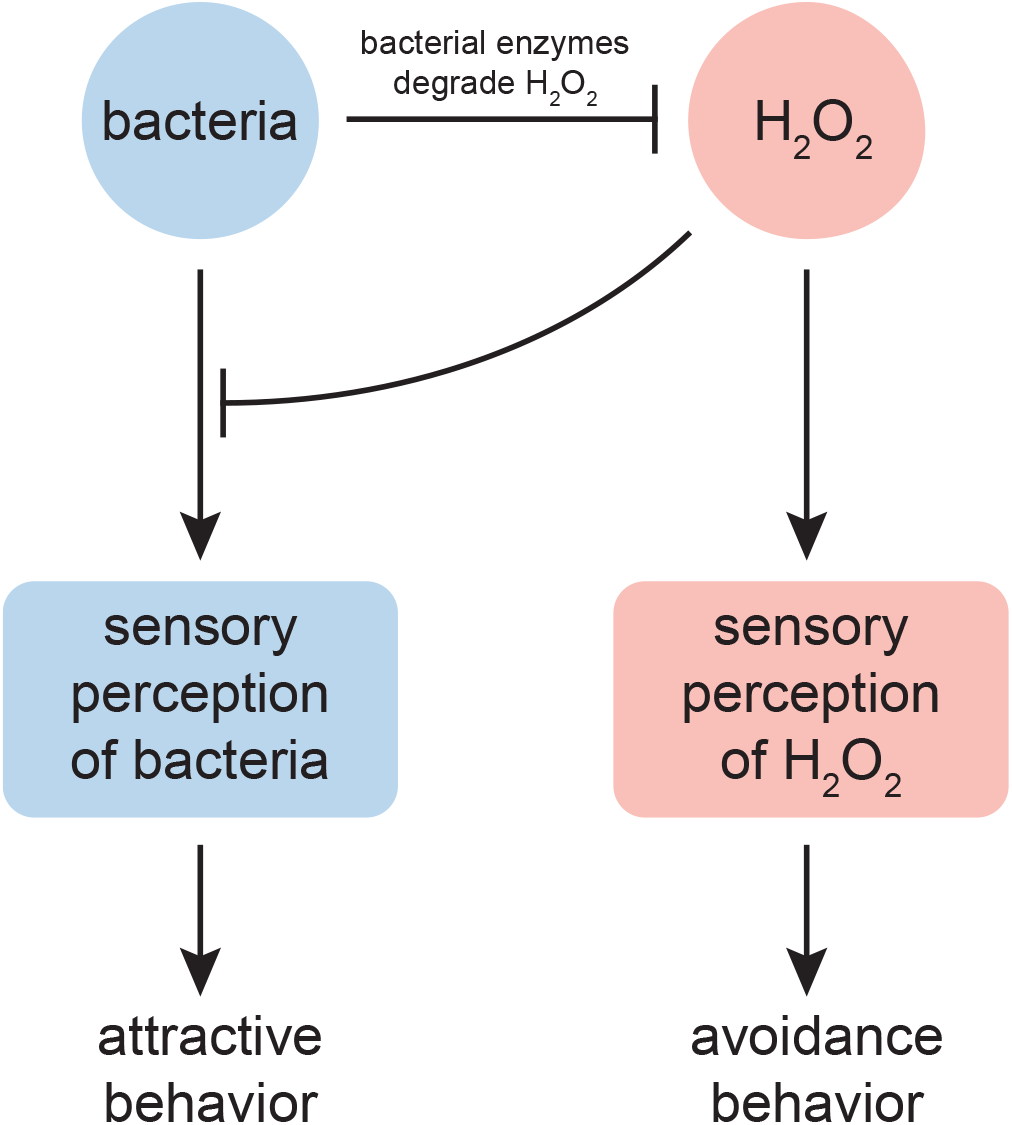
*C. elegans* assess faithfully the threat of hydrogen peroxide via sensory perception. H_2_O_2_ and bacteria trigger opposite locomotory behaviors through their action on the nematode’s sensory neurons. H_2_O_2_ and bacteria attenuate each other’s effects: H_2_O_2_ prevents or weakens sensory perception of bacteria and some bacteria express enzymes that degrade H_2_O_2_. The relative strength of this cross-inhibition leads to the differential sensory neuronal perception of H_2_O_2_ and bacteria, enabling the nematode to faithfully respond to the lethal threat of H_2_O_2_ by switching between locomotory behaviors that promote attraction or avoidance. H_2_O_2_ excites the ASJ sensory neurons that promote H_2_O_2_ avoidance. Bacteria excite, in a H_2_O_2_-sensitive manner, the ASEL, ADF, AWA, and BAG sensory neurons that promote attraction and inhibit the ASH and ASK neurons that promote aversion. The diminished sensory perception of bacteria unable to degrade H_2_O_2_ in the environment represents a general mechanism enabling nematodes to leave food patches that do not express sufficient levels of H_2_O_2_-degrading enzymes.

We show here that the ASJ neurons, which exhibited the largest response to H_2_O_2_, were necessary for *C. elegans* to escape environments with lethal H_2_O_2_ levels. This role is in line with ASJ’s well-established role mediating aversive responses to predator-secreted sulfolipids (Liu et al., 2018), pathogen secreted redox-active metabolites (Meisel et al., 2014), and short-wavelength light (Ward et al., 2008). The two other classes of sensory neurons that were excited by H_2_O_2_ are also known to mediate aversive responses: URX to high O_2_ levels (Chang et al., 2006; Cheung et al., 2005) and ASK to various water-soluble repellents (Gray et al., 2005; Hilliard et al., 2002). Previous studies showed that I2 sensory neurons in the *C. elegans* feeding organ are excited by H_2_O_2_ and inhibit bacterial ingestion in response to H_2_O_2_ (Bhatla and Horvitz, 2015) and olfactory neurons in the fruit fly *Drosophila* are excited by H_2_O_2_ produced in response to ultraviolet light and inhibit egg laying in response to that input (Guntur et al., 2017). We propose that excitation by H_2_O_2_ may be a common property of sensory neurons mediating aversive responses.

*C. elegans’* strong attraction to *E. coli* in the absence of environmental H_2_O_2_ was mirrored by the pattern of neuronal activity induced by *E. coli*. Exposure to *E. coli* excited sensory neurons that promote locomotion toward attractive cues (ASEL, ADF, AWA, and BAG) and generally inhibited sensory neurons that promote locomotion away from repulsive cues (ASH and ASK were inhibited, but AWB was excited). These findings generally follow and expand previous studies in *C. elegans* measuring neuronal modulation by *E. coli* (Calhoun et al., 2015; Iwanir et al., 2016) and by supernatants of *E. coli* culture medium (Ha et al., 2010; Wakabayashi et al., 2009; Zaslaver et al., 2015). The changes in sensory neuron activity induced by *E. coli* were prevented or weakened by environmental H_2_O_2_ when *E. coli* could not degrade H_2_O_2_. The diminished sensory perception of bacteria unable to degrade H_2_O_2_ in the environment represents a general mechanism enabling nematodes to leave food patches of laboratory and microbiome bacteria that do not express sufficient levels of H_2_O_2_-degrading enzymes.

We note that H_2_O_2_ prevented the responses to *E. coli* of a wide variety of sensory neurons. Previous studies in *C. elegans* showed that H_2_O_2_ blocks inactivating currents in ASER (Cai and Sesti, 2009) and that pretreatment with H_2_O_2_ blocked subsequent excitation of ASH sensory neurons by specific inputs (Li et al., 2016; Zhang et al., 2020) and lowered the spontaneous activity of AVA interneurons (Doser et al., 2020). The H_2_O_2_ concentration that we used in our studies is within the range produced by bacterial pathogens (Bolm et al., 2004; Jansen et al., 2002; Moy et al., 2004) and detected in inflammation or reperfusion after ischemia in mammals (Schroder and Eaton, 2008; Sprong et al., 1997). We propose that H_2_O_2_ may have a more widespread role in regulating neuronal activity than previously realized, and that physiological and pathological conditions may modulate neuronal activity by affecting H_2_O_2_ levels.

## Materials and Methods

### *C. elegans* culture, strains, and transgenes

Wild-type *C. elegans* was Bristol N2. *C. elegans* were cultured at 20°C on NGM agar plates (Nematode Growth Medium, 17 g/L agar, 2.5 g/L Bacto Peptone, 3.0 g/L NaCl, 1 mM CaCl_2_, 1 mM MgSO_4_, 25 mM H_2_KPO_4_/HK_2_PO4 pH 6.0, 5 mg/L cholesterol) seeded with *E. coli* OP50, unless noted otherwise. For a list of all bacterial and worm strains used in this study, see Tables S2, S3, and S4.

### Food choice assay

We performed two-choice food preference assays as described (Zhang et al., 2005), with some modifications. The assays were performed at a room temperature of 22°C on 10 cm chemotaxis agar plates (17 g/L agar, 1 mM CaCl_2_, 1 mM MgSO_4_, 25 mM H_2_KPO_4_/HK_2_PO4 pH 6.0). For assays with H_2_O_2_, we added the compound to the molten agar immediately before pouring. We added at opposite ends of each plate 50 µL of concentrated *E. coli* MG1655 and JI377 resuspended at an optical density (OD_600_) of 10 (Entchev et al., 2015), 20 hours before the beginning of the assay. We then transferred the agar from the petri plates and placed it on top of two glass slides about 2 inches apart on plastic containers filled half-way up the side of the agar with 1 mM H_2_O_2_ or water, as appropriate. This step ensured that bacteria on the plates could not deplete all the H_2_O_2_ on the plates if the compound were present. For assays with sodium azide, 5 µL of a 250 mM solution of the compound was added on top of each bacterial lawn 10 minutes before placing the worms. Day 1 adults derived from age-synchronized embryos obtained by bleaching were washed three times with M9 buffer, placed on the center of the agar, and the number of animals that reached each bacterial lawn was recorded at the specified times. The proportion of animals on a bacterial lawn was equal to (the number of animals on that lawn) / (total number of animals). The bacterial choice index was equal to (the number of animals on the MG1655 lawn - number of animals on the JI377 lawn) / (total number of animals).

### Food leaving assay

The assays were performed on 6 cm chemotaxis agar plates at a room temperature of 22°C. For assays with H_2_O_2_, we added the compound to the molten agar immediately before pouring. We added at the center of each plate 50 µL of concentrated bacteria resuspended at an OD_600_ of 10 (Entchev et al., 2015), 20 hours before the beginning of the assay. For assays where the JI377 lawn was derived from bacteria pre-treated with and without 1 mM H_2_O_2_, a bacterial resuspension in water at an OD_600_ of 10 was incubated at 20°C for 20 hours with or without 1 mM H_2_O_2_ and washed three times with water before plating. Unless noted, L4 larvae derived from age-synchronized embryos obtained by bleaching were washed three times with M9 buffer, placed on the center of the bacterial lawn, and the proportion of animals on the lawn was determined after two hours.

### H_2_O_2_ avoidance assay

We determined the proportion of aversive responses (avoidance index) as described previously (Liu et al., 2018). Briefly, we exposed day 1 adults on NGM plates without bacteria to a small volume of 1 mM H_2_O_2_ or water delivered with a glass capillary approximately 1 mm in front of the head of each animal. We tested each animal 5 times with each stimulus. An aversive response was defined as a reversal followed by an omega turn initiated within 4 seconds of exposure to the stimulus.

### H_2_O_2_ survival assays

We measured nematode development and survival on 6 cm chemotaxis agar plates with or without 1 mM H_2_O_2_, without bacteria or seeded with 50 µL *E. coli* MG1655 or JI377 resuspended at an OD_600_ of 10. For development assays, age-synchronized embryos obtained by bleaching were washed three times with M9 buffer, placed on the center of the bacterial lawn or plate, and their developmental stage was determined by visual inspection 72 hours later. For survival and reproduction assays, we transferred individual nematodes that had been in food leaving assays with JI377 and 1 mM H_2_O_2_ for 120 minutes to assay plates seeded with JI377 with or without 1 mM H_2_O_2_ added to the agar, and measured whether they died before reproduction, died after reproduction, or remained alive and reproduced after 72 hours. We also measured survival after adding 5 µL of MG1655 resuspended at OD_600_ of 10 to the center of the JI377 lawns of those food leaving assays.

### Behavioral state assay

We determined whether animals in food-leaving assays were roaming, dwelling, or quiescent by visual inspection, as described (Juozaityte et al., 2017). Briefly, animals were allowed to acclimate for 15 seconds, and the locomotion and pharyngeal pumping of each nematode was observed for 10 seconds using a dissection stereo microscope equipped with white-light transillumination. Roaming nematodes moved forward rapidly and turned infrequently, dwelling nematodes turned frequently or stayed in place while pumping, and quiescent nematodes were immobile and did not pump.

### Isolation and identification of compost microbiome bacteria

The MOYb collection of wild bacteria (Table S5) was isolated from nematodes isolated from residential compost in Massachusetts (USA). Small Tupperware containers containing manually homogenized compost were seeded with approximately 1000 eggs obtained by bleaching gravid adult N2, JU322, or PX178 worms, followed by three washes in S-basal media. Compost cultures were repeated in triplicate and incubated at room temperature ∼22ºC for 10 days. Following this culture, a small portion of compost was added to 10 cm NGM plates and worms were allowed to crawl onto the clean media, rinsed using M9 buffer and then plated onto clean NGM plates. Adult worms were then picked again to clean plates and allowed to crawl to spread bacterial colonies. This collection likely represents a combination of cuticle- and gut-associated bacteria. To ensure recovery of *C. elegans*, a mock culture was performed after which very few nematodes, largely of distinct morphology were recovered. Resulting bacterial colonies were isolated, grown on LB medium, and characterized via PCR using the indicated 16S rRNA gene primers (Table S5) 27f-YM AGAGTTTGATYMTGGCTCAG (Nercessian et al., 2005), 515f GTGCCAGCMGCCGCGGTAA (Turner et al., 1999), 806rB GGACTACNVGGGTWTCTAAT (Apprill et al., 2015), and 1492r GGTTACCTTGTTACGACTT (Turner et al., 1999). We performed phylogenetic analysis with CLC Main Workbench (Qiagen) based on partial 16S rRNA gene sequences using the neighbor-joining method and Jukes-Cantor correction. We placed age-synchronized embryos obtained by bleaching and washed three times with M9 buffer on bacterial lawns of specific MOYb strains on NGM plates and inspected those plates daily to determine whether the embryos grew to adulthood and reproduced.

### Sequence analyses of genes encoding hydrogen peroxide-degrading enzymes in microbiome bacterial genomes

We downloaded 180 sequenced bacterial genomes from the *C. elegans* microbiome (Zimmermann et al., 2020) along with *E. coli* MG1655 and *B. subtilis* PS832 using Joint Genome Institute’s IMG/M pipeline (https://img.jgi.doe.gov/) (Chen et al., 2021). Genome annotations were scanned for genes within clusters of orthologous groups (COGs) (Galperin et al., 2021) that have been associated with degradation of H_2_O_2_, including catalases, glutathione peroxidase, cytochrome c peroxidase, peroxiredoxins, and rubrerythrins. We performed additional protein alignments of KatE and KatG orthologs using ClustalOmega (Sievers et al., 2011). The resulting alignments were analyzed through SnapGene (Version 5.3.2) for the presence of key catalytic or ligand binding residues noted by the Protein Data Bank in Europe - Knowledge Base (PDBe-KB).

### Catalase assays

We screened the catalase activity of bacterial strains in specific *C. elegans* microbiome collections by observing the extent to which oxygen bubbles were formed after mixing on a glass slide a drop of 9.8 M H_2_O_2_ with an inoculation loop loaded with a sample of a bacterial colony grown on a LB plate at room temperature. Quantitative catalase assays were performed as described (Iwase et al., 2013). Briefly, catalase activity was measured as the height of the foam column of oxygen bubbles formed in a test tube with 0.1 mL of 9.8 M H_2_O_2_, 0.1 mL of 1% Triton X-100, and 0.1 mL of a bacterial culture in LB broth resuspended at an OD_600_ of 10. We generated full bacterial growth curves to identify when bacteria were in exponential growth and stationary phase.

### Calcium imaging

We picked L4 stage worms and, within six hours, placed them in a microfluidic device that allowed delivery of stimuli alternating with buffer (Si et al., 2019). We delivered each stimulus for 15 seconds, separated by buffer (H_2_O) for 45 seconds. Each animal was presented twice with the set of 6 stimuli. The order of stimuli was randomized on each delivery to control for possible temporal interactions. The stimuli consisted of water, or *E. coli* MG1655 or JI377 cultured overnight in LB broth at 37°C and then resuspended in water at an OD_600_ of 10, each incubated at 20°C for 20 hours with or without 1 mM H_2_O_2_. Fluorescence was recorded with a spinning disc confocal microscope (Dragonfly 200, Andor) and a sCMOS camera (Andor Zyla 4.2p) that captured fluorescence from GCaMP6s at 10 ms/1.2 µm z-slice, 25 z-slices/volume, and 4 volumes/second. To extract calcium activity from the recorded data, we identified the center of each neuronal nucleus in every frame and took the average pixel intensities of a 3.2 µm x 3.2 µm x 3.6 µm rectangular box around those centers. The neuron-independent background signal was removed and ΔF/F_0_ calculated for each stimulus-response, where F_0_ was the average fluorescence value during the five seconds before delivery of the stimulus. We used Morpheus (https://software.broadinstitute.org/morpheus/) to perform two-way hierarchical clustering of mean ΔF/F_0_ values (at the 0.5 second period at the center of each stimulus interval) of the 26 sensory neurons across the 6 stimuli.

### Statistical analysis

Statistical analyses were performed in JMP Pro version 15 (SAS). We tested for differences in the average of chemotaxis, food choice, food leaving, H_2_O_2_ avoidance, and GCaMP6 fluorescence using ANOVA. We used the Tukey HSD post-hoc test to determine which pairs of groups in the sample differ, in cases where more than two groups were compared. We used Fisher’s exact test to determine whether the proportion of dead animals were equal across groups. We used ordinal and least-squares regression to quantify interactions between groups using the following linear model: data = Intercept + group 1 + group 2 + group 1 * group 2*+ ε*. The second to last term in this model quantifies the existence, magnitude, and type (synergistic or antagonistic) of interaction between groups. We used the Bonferroni correction to adjust *P* values when performing multiple comparisons.

## Acknowledgements

We thank Erin Cram, Edward Geisinger, and Julian Stanley for detailed comments on our manuscript. Joy Alcedo, James Imlay, Dennis Kim, Gary Ruvkun, Piali Sengupta, and Peter Setlow kindly provided strains. We benefitted from discussions with members of Javier Apfeld’s and Erin Cram’s labs, Yunrong Chai, Marie-Anne Felix, and Deborah Gordon. Some strains were provided by the CGC, which is funded by NIH Office of Research Infrastructure Programs (P40 OD010440). The research was supported by National Science Foundation CAREER grant #1750065 to J.A., Burroughs Wellcome Fund and American Federation for Aging Research awards to V.V., NIH grant DP2DK116645 to B.S.S., JGI/DOE grant CSP503338 to B.S.S., NSF DBI Research Experiences for Undergraduates Award #1757443 to O.B., and Northeastern University Tier 1 award to V.V. and J.A,

