## Supplementary information for "Modulation of sensory perception by hydrogen peroxide enables Caenorhabditis elegans to find a niche that provides both food and protection from hydrogen peroxide"

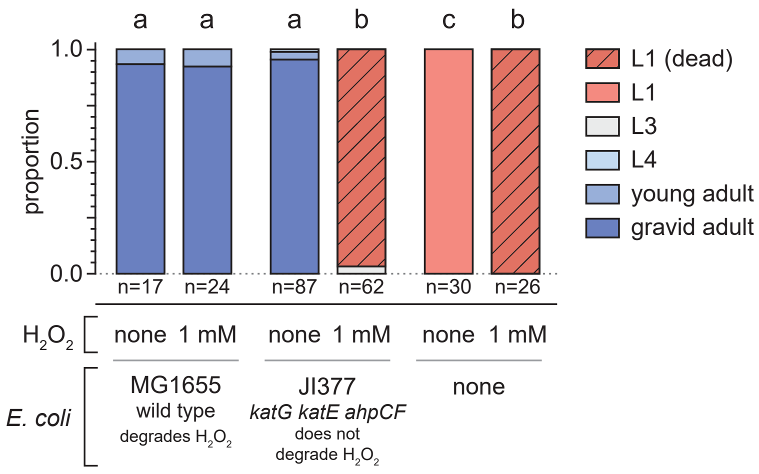


**Supplementary Figure 1. $H_2O_2$-degrading enzymes from *E. coli* create an environment where *C. elegans* is safe from the threat of $H_2O_2$**

Development of wild-type *C. elegans* embryos in the presence of 1 mM $H_2O_2$. Groups labeled with different letters exhibited significant differences ($P < 0.001$, ordinal logistic regression) otherwise ($P > 0.05$).

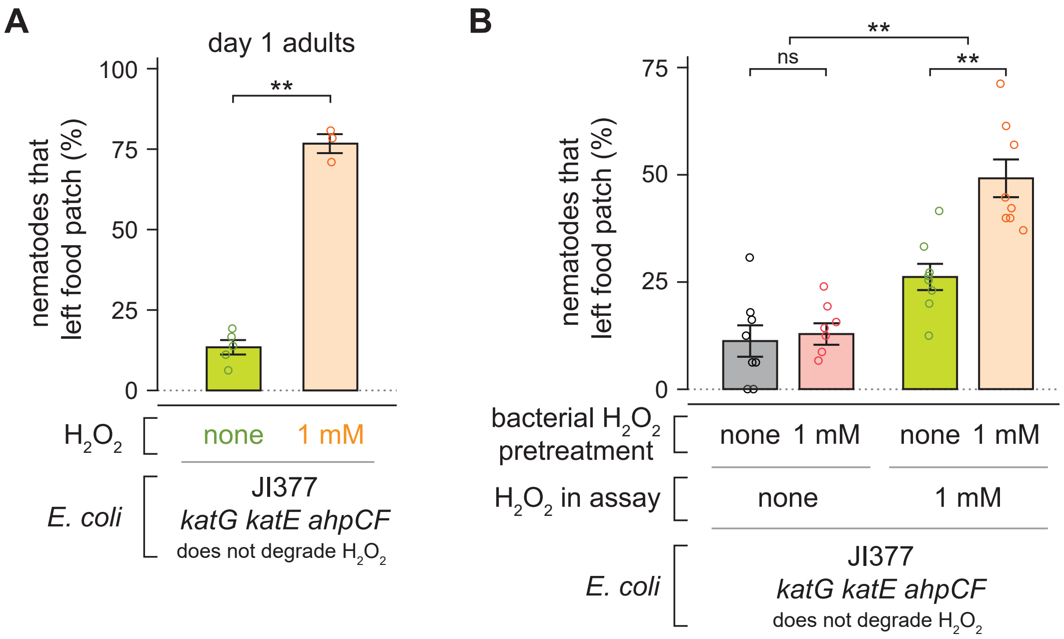


**Supplementary Figure 2. Hydrogen peroxide induces food leaving in adult *C. elegans***

(A) $H_2O_2$ induced an increase in the proportion of day 1 adult *C. elegans* nematodes that left a patch of *E. coli* JI377. ** indicates $P < 0.0001$ (t-test).
(B) Pre-treating the *E. coli* JI377 suspension used to make the lawn with 1 mM $H_2O_2$ for 20 hours did not increase nematode lawn leaving when no $H_2O_2$ was added to the assay plates, but caused a larger increase in lawn leaving when 1 mM $H_2O_2$ was added to the plates. ** indicates $P < 0.006$ and "ns" indicates $P > 0.05$ (standard least-squares regression).
Data are represented as mean ± s.e.m.

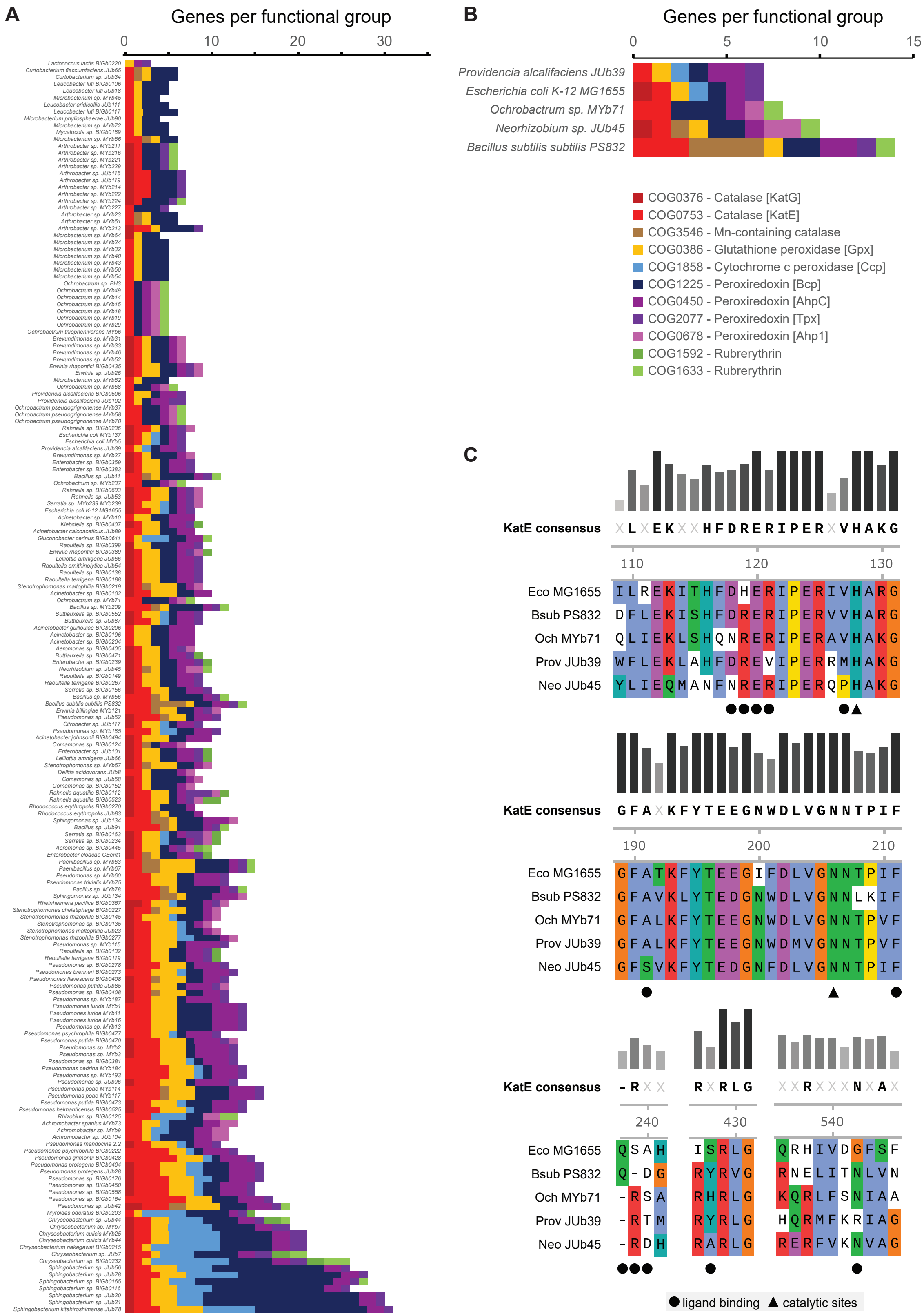


**Supplementary Figure 3. Distribution and alignments of hydrogen peroxide degrading enzymes in sequenced genomes of *C. elegans* natural microbiome.**

(A-B) We identified any genes within clusters of orthologous groups (COGs) associated with $H_2O_2$

degrading capabilities in 180 sequenced *C. elegans* microbiome genomes, plus *E. coli* MG1655 and *B. subtilis* PS832. These include catalase (COG0376, COG0753, COG3546), glutathione peroxidase (COG0386), cytochrome c peroxidase (COG1858), peroxiredoxins (COG1225, COG0450, COG2077, COG0678) and rubrerythrins (COG1592, COG1633). Total genes within each class are plotted for each strain

(C) Selected regions of protein alignments for a subset of the KatE orthologs (COG0753) are highlighted to show conservation of the catalytic residues (triangles) and variation in the predicted $H_2O_2$ ligand binding residues (based on *E. coli*).

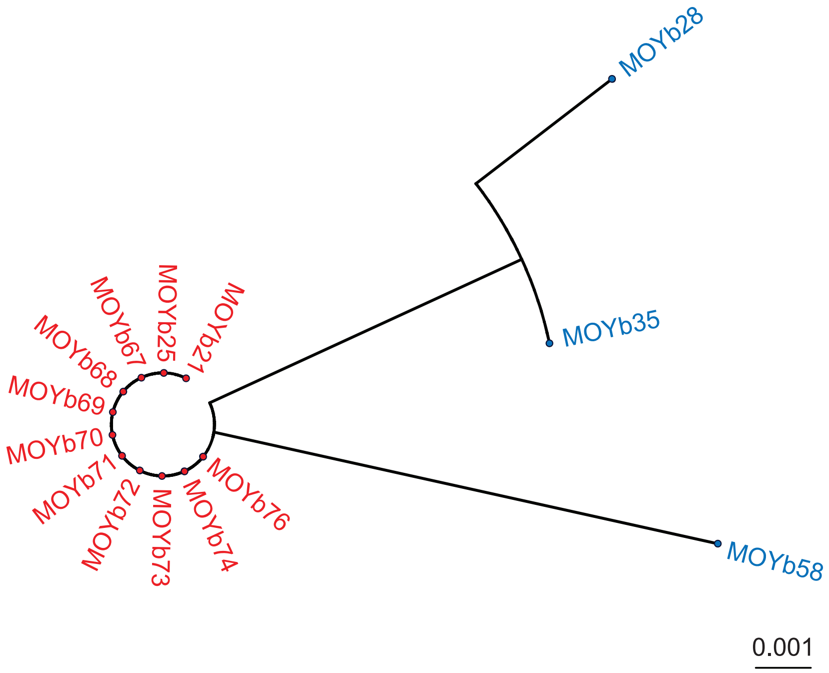


**Supplementary Figure 4. Phylogeny of *Shewanella* strains from compost microbiome**

Phylogenetic tree, reconstructed using the neighbor-joining method based on partial 16S rRNA gene sequences, indicating the relationships among 14 *Shewanella* strains isolated from compost microbiome. The scale bar indicates the number of substitutions per site. *Shewanella* strains that supported *C. elegans* growth and reproduction are denoted in blue and strains that did not and were, instead, pathogenic to *C. elegans* are denoted in red. The isolation notes for these bacterial strains are shown in Table S5.

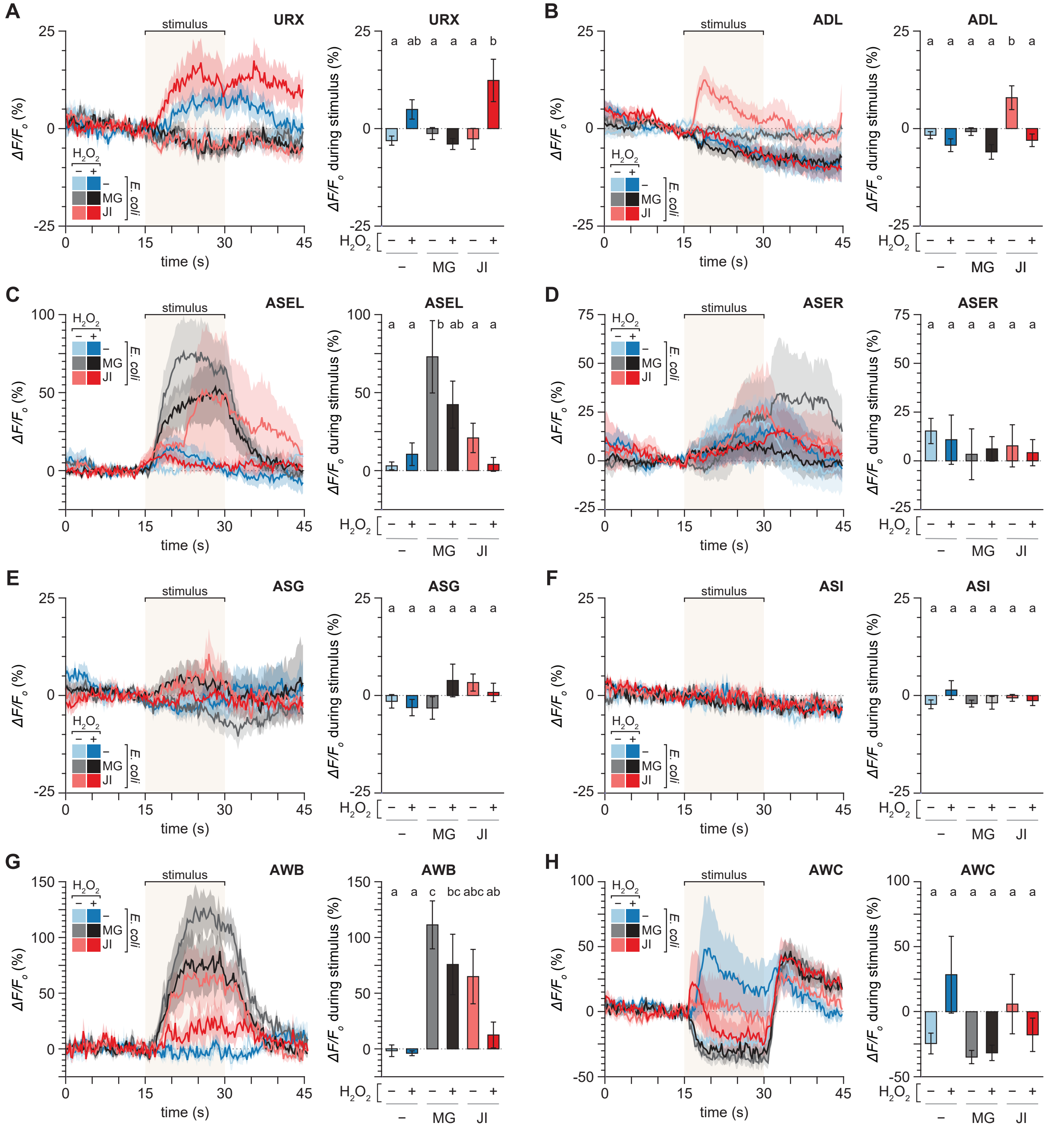


**Supplementary Figure 5. Hydrogen peroxide and bacteria have opposing effects on the activity of sensory neurons**

(A-H) Average GCaMP6 fluorescence traces of (A) URX, (B) ADL, (C) ASEL, (D) ASER, (E) ASG, (F) ASI, (G) AWB, and (H) AWC neuronal classes in response to six different stimuli (left sub-panels) and average changes in fluorescence in response to those stimuli (right sub-panels). The stimulus delivery interval is indicated by a shaded box. Data are represented as mean ± s.e.m. The number of neurons imaged was 28 ADF, 28 ADL, 14 ASEL, 14 ASER, 28 ASG, 28 ASH, 27 ASI, 28 ASJ, 28 ASK, 28 AWA, 13 AWB, 24 AWC, 18 BAG, and 28 URX. Groups labeled with different letters exhibited significant differences ($P < 0.05$, Tukey HSD test) otherwise ($P > 0.05$). Traces for the ASJ, ADF, AWA, BAG, ASK, and ASH neuronal classes are shown in Figure 4.

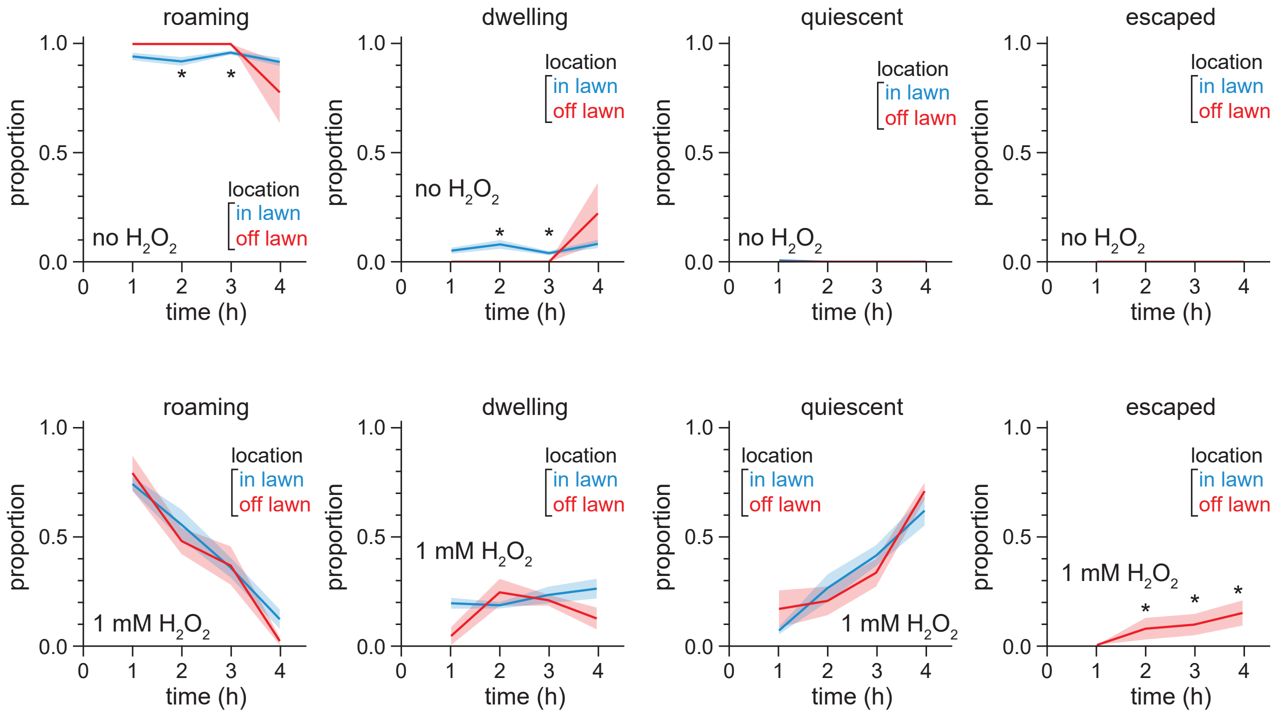


**Supplementary Figure 6. Nematode behavioral state in response to hydrogen peroxide**

The plots show the proportion of animals roaming, dwelling, and quiescent that stayed in the *E. coli* JI377 lawn (blue) or left the lawn (red), and the proportion of animals that escaped the plate after leaving the lawn (red), in the assays on plates without $H_2O_2$ (top row) and on plates with 1 mM $H_2O_2$ (bottom row). Data are represented as mean ± s.e.m of the n = 6 assays per condition shown in Figure 6B. * indicates $P < 0.05$ otherwise $P > 0.05$ (t-test).

**Supplementary Table 1. Genes encoding hydrogen peroxide-degrading enzymes in sequenced bacterial strains isolated from *C. elegans* natural microbiome.**

| Genome | COG0376 - Catalase [KatG] | COG0753 - Catalase [KatE] | COG3546 - Mn-containing catalase | COG0386 - Glutathione peroxidase [Gpx] | COG1858 - Cytochrome c peroxidase [Ccp] | COG0450 - Peroxiredoxin [AhpC] | COG1225 - Peroxiredoxin [Bcp] | COG2077 - Peroxiredoxin [Tpx] | COG0678 - Peroxiredoxin [Ahp1] | COG1592 - Rubrerythrin | COG1633 - Rubrerythrin | Total |
|---|---|---|---|---|---|---|---|---|---|---|---|---|
| Lactococcus lactis BIGb0220 | 0 | 0 | 0 | 1 | 0 | 1 | 0 | 0 | 1 | 0 | 0 | 3 |
| Microbacterium phyllosphaerae JUb90 | 0 | 1 | 0 | 1 | 0 | 0 | 0 | 2 | 0 | 0 | 0 | 4 |
| Microbacterium sp. MYb45 | 0 | 1 | 0 | 1 | 0 | 0 | 0 | 2 | 0 | 0 | 0 | 4 |
| Microbacterium sp. MYb64 | 1 | 0 | 0 | 1 | 0 | 0 | 0 | 2 | 0 | 0 | 0 | 4 |
| Arthrobacter sp. MYb227 | 0 | 1 | 0 | 0 | 0 | 0 | 0 | 3 | 1 | 0 | 0 | 5 |
| Leucobacter aridicollis JUb111 | 0 | 1 | 0 | 1 | 0 | 0 | 0 | 3 | 0 | 0 | 0 | 5 |
| Leucobacter luti BIGb0106 | 0 | 1 | 0 | 1 | 0 | 0 | 0 | 3 | 0 | 0 | 0 | 5 |
| Leucobacter luti JUb18 | 0 | 1 | 0 | 1 | 0 | 0 | 0 | 3 | 0 | 0 | 0 | 5 |
| Microbacterium sp. MYb24 | 0 | 1 | 0 | 1 | 0 | 0 | 0 | 3 | 0 | 0 | 0 | 5 |
| Microbacterium sp. MYb32 | 0 | 1 | 0 | 1 | 0 | 0 | 0 | 3 | 0 | 0 | 0 | 5 |
| Microbacterium sp. MYb40 | 0 | 1 | 0 | 1 | 0 | 0 | 0 | 3 | 0 | 0 | 0 | 5 |
| Microbacterium sp. MYb43 | 0 | 1 | 0 | 1 | 0 | 0 | 0 | 3 | 0 | 0 | 0 | 5 |
| Microbacterium sp. MYb50 | 0 | 1 | 0 | 1 | 0 | 0 | 0 | 3 | 0 | 0 | 0 | 5 |
| Microbacterium sp. MYb54 | 0 | 1 | 0 | 1 | 0 | 0 | 0 | 3 | 0 | 0 | 0 | 5 |
| Microbacterium sp. MYb62 | 0 | 1 | 0 | 1 | 0 | 0 | 0 | 3 | 0 | 0 | 0 | 5 |
| Microbacterium sp. MYb72 | 0 | 1 | 0 | 1 | 0 | 0 | 0 | 3 | 0 | 0 | 0 | 5 |
| Mycetocola sp. BIGb0189 | 0 | 1 | 0 | 1 | 0 | 0 | 0 | 3 | 0 | 0 | 0 | 5 |
| Ochrobactrum sp. BH3 | 0 | 1 | 0 | 0 | 0 | 1 | 1 | 1 | 0 | 0 | 1 | 5 |
| Ochrobactrum sp. MYb14 | 0 | 1 | 0 | 0 | 0 | 1 | 1 | 1 | 0 | 0 | 1 | 5 |
| Ochrobactrum sp. MYb15 | 0 | 1 | 0 | 0 | 0 | 1 | 1 | 1 | 0 | 0 | 1 | 5 |
| Ochrobactrum sp. MYb18 | 0 | 1 | 0 | 0 | 0 | 1 | 1 | 1 | 0 | 0 | 1 | 5 |
| Ochrobactrum sp. MYb19 | 0 | 1 | 0 | 0 | 0 | 1 | 1 | 1 | 0 | 0 | 1 | 5 |
| Ochrobactrum sp. MYb29 | 0 | 1 | 0 | 0 | 0 | 1 | 1 | 1 | 0 | 0 | 1 | 5 |
| Ochrobactrum sp. MYb49 | 0 | 1 | 0 | 0 | 0 | 1 | 1 | 1 | 0 | 0 | 1 | 5 |
| Ochrobactrum thiophenivorans MYb6 | 0 | 1 | 0 | 0 | 0 | 1 | 1 | 1 | 0 | 0 | 1 | 5 |
| Arthrobacter sp. MYb211 | 1 | 1 | 0 | 0 | 0 | 0 | 0 | 2 | 1 | 0 | 1 | 6 |
| Arthrobacter sp. MYb216 | 1 | 1 | 0 | 0 | 0 | 0 | 0 | 2 | 1 | 0 | 1 | 6 |
| Arthrobacter sp. MYb221 | 1 | 1 | 0 | 0 | 0 | 0 | 0 | 2 | 1 | 0 | 1 | 6 |
| Arthrobacter sp. MYb229 | 1 | 1 | 0 | 0 | 0 | 0 | 0 | 2 | 1 | 0 | 1 | 6 |
| Arthrobacter sp. MYb23 | 0 | 1 | 1 | 1 | 0 | 0 | 0 | 3 | 0 | 0 | 0 | 6 |
| Arthrobacter sp. MYb51 | 0 | 1 | 1 | 1 | 0 | 0 | 0 | 3 | 0 | 0 | 0 | 6 |
| Curtobacterium flaccumfaciens JUb65 | 0 | 1 | 1 | 1 | 0 | 0 | 0 | 3 | 0 | 0 | 0 | 6 |
| Curtobacterium sp. JUb34 | 0 | 1 | 1 | 1 | 0 | 0 | 0 | 3 | 0 | 0 | 0 | 6 |
| Leucobacter luti BIGb0117 | 0 | 1 | 0 | 1 | 0 | 0 | 0 | 4 | 0 | 0 | 0 | 6 |
| Microbacterium sp. MYb66 | 0 | 2 | 1 | 1 | 0 | 0 | 0 | 2 | 0 | 0 | 0 | 6 |
| Ochrobactrum sp. MYb68 | 0 | 1 | 0 | 0 | 0 | 1 | 1 | 2 | 0 | 0 | 1 | 6 |
| Arthrobacter sp. JUb115 | 1 | 2 | 0 | 0 | 0 | 0 | 0 | 3 | 1 | 0 | 0 | 7 |
| Arthrobacter sp. JUb119 | 1 | 2 | 0 | 0 | 0 | 0 | 0 | 3 | 1 | 0 | 0 | 7 |
| Arthrobacter sp. MYb214 | 1 | 2 | 0 | 0 | 0 | 0 | 0 | 3 | 1 | 0 | 0 | 7 |
| Arthrobacter sp. MYb222 | 1 | 2 | 0 | 0 | 0 | 0 | 0 | 3 | 1 | 0 | 0 | 7 |
| Arthrobacter sp. MYb224 | 1 | 1 | 0 | 0 | 0 | 0 | 0 | 3 | 1 | 0 | 1 | 7 |
| Brevundimonas sp. MYb31 | 1 | 1 | 0 | 1 | 0 | 1 | 1 | 2 | 0 | 0 | 0 | 7 |
| Brevundimonas sp. MYb33 | 1 | 1 | 0 | 1 | 0 | 1 | 1 | 2 | 0 | 0 | 0 | 7 |
| Brevundimonas sp. MYb46 | 1 | 1 | 0 | 1 | 0 | 1 | 1 | 2 | 0 | 0 | 0 | 7 |
| Brevundimonas sp. MYb52 | 1 | 1 | 0 | 1 | 0 | 1 | 1 | 2 | 0 | 0 | 0 | 7 |
| Comamonas sp. BIGb0124 | 1 | 1 | 1 | 0 | 0 | 2 | 1 | 1 | 0 | 0 | 0 | 7 |
| Escherichia coli K-12 MG1655 | 1 | 1 | 0 | 1 | 1 | 1 | 0 | 1 | 1 | 0 | 0 | 7 |

| Genome | COG0376 - Catalase [KatG] | COG0753 - Catalase [KatE] | COG3546 - Mn-containing catalase | COG0386 - Glutathione peroxidase [Gpx] | COG1858 - Cytochrome c peroxidase [Ccp] | COG0450 - Peroxiredoxin [AhpC] | COG1225 - Peroxiredoxin [Bcp] | COG2077 - Peroxiredoxin [Tpx] | COG0678 - Peroxiredoxin [Ahp1] | COG1592 - Rubrerythrin | COG1633 - Rubrerythrin | Total |
|---|---|---|---|---|---|---|---|---|---|---|---|---|
| Escherichia coli MYb137 | 1 | 1 | 0 | 1 | 1 | 1 | 0 | 1 | 1 | 0 | 0 | 7 |
| Escherichia coli MYb5 | 1 | 1 | 0 | 1 | 1 | 1 | 0 | 1 | 1 | 0 | 0 | 7 |
| Ochrobactrum pseudogrignonense MYb37 | 0 | 2 | 0 | 0 | 0 | 1 | 1 | 2 | 0 | 0 | 1 | 7 |
| Ochrobactrum pseudogrignonense MYb58 | 0 | 2 | 0 | 0 | 0 | 1 | 1 | 2 | 0 | 0 | 1 | 7 |
| Ochrobactrum pseudogrignonense MYb70 | 0 | 2 | 0 | 0 | 0 | 1 | 1 | 2 | 0 | 0 | 1 | 7 |
| Ochrobactrum sp. MYb237 | 0 | 2 | 0 | 0 | 0 | 1 | 1 | 2 | 0 | 0 | 1 | 7 |
| Providencia alcalifaciens BIGb0506 | 0 | 1 | 0 | 2 | 0 | 2 | 0 | 1 | 1 | 0 | 0 | 7 |
| Providencia alcalifaciens JUb102 | 0 | 1 | 0 | 1 | 0 | 2 | 0 | 1 | 2 | 0 | 0 | 7 |
| Providencia alcalifaciens JUb39 | 0 | 1 | 0 | 1 | 1 | 2 | 0 | 1 | 1 | 0 | 0 | 7 |
| Acinetobacter calcoaceticus JUb89 | 0 | 2 | 0 | 2 | 0 | 2 | 0 | 2 | 0 | 0 | 0 | 8 |
| Acinetobacter guillouiae BIGb0206 | 0 | 1 | 0 | 2 | 0 | 3 | 0 | 2 | 0 | 0 | 0 | 8 |
| Acinetobacter sp. BIGb0196 | 0 | 1 | 0 | 2 | 0 | 3 | 0 | 2 | 0 | 0 | 0 | 8 |
| Acinetobacter sp. BIGb0204 | 0 | 1 | 0 | 2 | 0 | 3 | 0 | 2 | 0 | 0 | 0 | 8 |
| Acinetobacter sp. MYb10 | 0 | 1 | 0 | 2 | 0 | 3 | 0 | 2 | 0 | 0 | 0 | 8 |
| Aeromonas sp. BIGb0405 | 1 | 1 | 0 | 2 | 0 | 1 | 1 | 1 | 1 | 0 | 0 | 8 |
| Brevundimonas sp. MYb27 | 1 | 1 | 0 | 1 | 0 | 1 | 1 | 3 | 0 | 0 | 0 | 8 |
| Comamonas sp. BIGb0152 | 1 | 1 | 0 | 1 | 0 | 1 | 1 | 3 | 0 | 0 | 0 | 8 |
| Delftia acidovorans JUb8 | 0 | 2 | 0 | 1 | 0 | 1 | 1 | 3 | 0 | 0 | 0 | 8 |
| Enterobacter sp. BIGb0359 | 1 | 1 | 0 | 2 | 0 | 2 | 0 | 1 | 1 | 0 | 0 | 8 |
| Enterobacter sp. BIGb0383 | 1 | 1 | 0 | 2 | 0 | 2 | 0 | 1 | 1 | 0 | 0 | 8 |
| Ochrobactrum sp. MYb71 | 0 | 2 | 0 | 0 | 0 | 1 | 1 | 3 | 0 | 0 | 1 | 8 |
| Rahnella sp. BIGb0236 | 1 | 2 | 0 | 1 | 0 | 1 | 1 | 1 | 1 | 0 | 0 | 8 |
| Rhodococcus erythropolis BIGb0270 | 1 | 2 | 0 | 1 | 0 | 1 | 0 | 3 | 0 | 0 | 0 | 8 |
| Arthrobacter sp. MYb213 | 1 | 2 | 0 | 1 | 0 | 0 | 0 | 4 | 1 | 0 | 0 | 9 |
| Buttiauxella sp. BIGb0471 | 1 | 1 | 0 | 2 | 0 | 2 | 0 | 1 | 1 | 1 | 0 | 9 |
| Buttiauxella sp. BIGb0552 | 1 | 1 | 0 | 2 | 1 | 2 | 0 | 1 | 1 | 0 | 0 | 9 |
| Buttiauxella sp. JUb87 | 1 | 2 | 0 | 1 | 1 | 2 | 0 | 1 | 1 | 0 | 0 | 9 |
| Citrobacter sp. JUb117 | 1 | 1 | 0 | 2 | 1 | 2 | 0 | 1 | 1 | 0 | 0 | 9 |
| Comamonas sp. JUb58 | 1 | 1 | 0 | 1 | 0 | 1 | 1 | 4 | 0 | 0 | 0 | 9 |
| Erwinia rhapontici BIGb0435 | 1 | 1 | 0 | 2 | 0 | 1 | 1 | 2 | 1 | 0 | 0 | 9 |
| Erwinia sp. JUb26 | 1 | 1 | 0 | 2 | 0 | 1 | 1 | 2 | 1 | 0 | 0 | 9 |
| Klebsiella sp. BIGb0407 | 1 | 1 | 0 | 2 | 0 | 2 | 0 | 1 | 1 | 1 | 0 | 9 |
| Lelliottia amnigena JUb66 | 1 | 1 | 0 | 2 | 0 | 2 | 0 | 2 | 1 | 0 | 0 | 9 |
| Rahnella sp. BIGb0603 | 1 | 2 | 0 | 2 | 0 | 1 | 1 | 1 | 1 | 0 | 0 | 9 |
| Rahnella sp. JUb53 | 1 | 2 | 0 | 2 | 0 | 1 | 1 | 1 | 1 | 0 | 0 | 9 |
| Raoultella ornithinolytica JUb54 | 1 | 1 | 0 | 2 | 0 | 2 | 0 | 2 | 1 | 0 | 0 | 9 |
| Raoultella sp. BIGb0138 | 1 | 1 | 0 | 2 | 0 | 2 | 0 | 2 | 1 | 0 | 0 | 9 |
| Raoultella sp. BIGb0149 | 1 | 1 | 0 | 2 | 0 | 2 | 0 | 2 | 1 | 0 | 0 | 9 |
| Raoultella sp. BIGb0399 | 1 | 0 | 0 | 2 | 0 | 2 | 0 | 3 | 1 | 0 | 0 | 9 |
| Raoultella terrigena BIGb0188 | 1 | 1 | 0 | 2 | 0 | 2 | 0 | 2 | 1 | 0 | 0 | 9 |
| Raoultella terrigena BIGb0267 | 1 | 1 | 0 | 2 | 0 | 2 | 0 | 2 | 1 | 0 | 0 | 9 |
| Rhodococcus erythropolis JUb83 | 1 | 2 | 1 | 1 | 0 | 1 | 0 | 3 | 0 | 0 | 0 | 9 |
| Serratia sp. MYb239 MYb239 | 1 | 1 | 0 | 2 | 1 | 1 | 1 | 1 | 1 | 0 | 0 | 9 |
| Stenotrophomonas maltophilia BIGb0219 | 1 | 1 | 1 | 3 | 0 | 1 | 1 | 1 | 0 | 0 | 0 | 9 |
| Acinetobacter johnsonii BIGb0494 | 1 | 1 | 0 | 3 | 0 | 3 | 0 | 2 | 0 | 0 | 0 | 10 |
| Acinetobacter sp. BIGb0102 | 1 | 2 | 0 | 3 | 0 | 2 | 0 | 2 | 0 | 0 | 0 | 10 |
| Aeromonas sp. BIGb0445 | 1 | 1 | 0 | 2 | 0 | 1 | 1 | 1 | 1 | 1 | 1 | 10 |

| Genome | COG0376 - Catalase [KatG] | COG0753 - Catalase [KatE] | COG3546 - Mn-containing catalase | COG0386 - Glutathione peroxidase [Gpx] | COG1858 - Cytochrome c peroxidase [Ccp] | COG0450 - Peroxiredoxin [AhpC] | COG1225 - Peroxiredoxin [Bcp] | COG2077 - Peroxiredoxin [Tpx] | COG0678 - Peroxiredoxin [Ahp1] | COG1592 - Rubrerythrin | COG1633 - Rubrerythrin | Total |
|---|---|---|---|---|---|---|---|---|---|---|---|---|
| Enterobacter cloacae CEent1 | 1 | 1 | 1 | 1 | 0 | 2 | 0 | 2 | 1 | 1 | 0 | 10 |
| Enterobacter sp. BIGb0239 | 1 | 1 | 0 | 2 | 0 | 2 | 0 | 2 | 1 | 1 | 0 | 10 |
| Enterobacter sp. JUb101 | 1 | 1 | 0 | 2 | 0 | 2 | 0 | 2 | 1 | 1 | 0 | 10 |
| Erwinia rhapontici BIGb0389 | 1 | 1 | 0 | 2 | 0 | 1 | 1 | 2 | 1 | 1 | 0 | 10 |
| Gluconobacter cerinus BIGb0611 | 0 | 1 | 0 | 1 | 3 | 2 | 0 | 2 | 0 | 0 | 1 | 10 |
| Lelliottia amnigena JUb66 | 1 | 1 | 0 | 2 | 0 | 2 | 0 | 2 | 1 | 1 | 0 | 10 |
| Neorhizobium sp. JUb45 | 1 | 1 | 1 | 1 | 0 | 1 | 2 | 2 | 0 | 0 | 1 | 10 |
| Serratia sp. BIGb0156 | 1 | 2 | 0 | 2 | 0 | 2 | 0 | 2 | 1 | 0 | 0 | 10 |
| Stenotrophomonas sp. BIGb0135 | 1 | 2 | 0 | 3 | 1 | 1 | 1 | 1 | 0 | 0 | 0 | 10 |
| Stenotrophomonas sp. MYb57 | 1 | 1 | 1 | 3 | 0 | 1 | 1 | 2 | 0 | 0 | 0 | 10 |
| Bacillus sp. JUb11 | 0 | 2 | 1 | 1 | 0 | 1 | 0 | 4 | 1 | 0 | 1 | 11 |
| Erwinia billingiae MYb121 | 0 | 3 | 1 | 2 | 0 | 1 | 1 | 2 | 1 | 0 | 0 | 11 |
| Pseudomonas mendocina 2.2 | 1 | 2 | 0 | 3 | 1 | 2 | 0 | 1 | 1 | 0 | 0 | 11 |
| Pseudomonas psychrophila BIGb0477 | 0 | 3 | 0 | 2 | 1 | 3 | 0 | 1 | 1 | 0 | 0 | 11 |
| Pseudomonas sp. JUb52 | 1 | 3 | 1 | 2 | 0 | 2 | 0 | 1 | 1 | 0 | 0 | 11 |
| Pseudomonas sp. MYb185 | 0 | 3 | 0 | 1 | 2 | 2 | 0 | 2 | 1 | 0 | 0 | 11 |
| Rahnella aquatilis BIGb0523 | 1 | 2 | 0 | 2 | 0 | 1 | 1 | 1 | 1 | 2 | 0 | 11 |
| Raoultella sp. BIGb0132 | 1 | 2 | 0 | 2 | 0 | 2 | 0 | 2 | 1 | 1 | 0 | 11 |
| Raoultella terrigena BIGb0119 | 1 | 2 | 0 | 2 | 0 | 2 | 0 | 2 | 1 | 1 | 0 | 11 |
| Serratia sp. BIGb0163 | 1 | 1 | 0 | 2 | 1 | 1 | 1 | 1 | 2 | 1 | 0 | 11 |
| Serratia sp. BIGb0234 | 1 | 1 | 0 | 2 | 1 | 1 | 1 | 2 | 1 | 1 | 0 | 11 |
| Stenotrophomonas chelatiphaga BIGb0227 | 1 | 2 | 1 | 3 | 0 | 1 | 1 | 2 | 0 | 0 | 0 | 11 |
| Stenotrophomonas maltophilia JUb23 | 1 | 2 | 0 | 3 | 1 | 1 | 1 | 2 | 0 | 0 | 0 | 11 |
| Stenotrophomonas rhizophila BIGb0145 | 1 | 1 | 0 | 3 | 1 | 1 | 1 | 3 | 0 | 0 | 0 | 11 |
| Achromobacter sp. JUb104 | 0 | 2 | 0 | 1 | 1 | 2 | 2 | 4 | 0 | 0 | 0 | 12 |
| Achromobacter sp. MYb9 | 0 | 2 | 0 | 1 | 2 | 2 | 2 | 3 | 0 | 0 | 0 | 12 |
| Bacillus sp. JUb91 | 0 | 4 | 1 | 1 | 0 | 1 | 0 | 3 | 1 | 0 | 1 | 12 |
| Bacillus sp. MYb209 | 0 | 3 | 1 | 1 | 0 | 1 | 0 | 4 | 1 | 0 | 1 | 12 |
| Bacillus sp. MYb56 | 0 | 3 | 1 | 1 | 0 | 1 | 0 | 4 | 1 | 0 | 1 | 12 |
| Pseudomonas flavescens BIGb0408 | 1 | 2 | 1 | 3 | 0 | 2 | 0 | 2 | 1 | 0 | 0 | 12 |
| Pseudomonas putida JUb85 | 1 | 2 | 0 | 3 | 0 | 3 | 1 | 1 | 1 | 0 | 0 | 12 |
| Pseudomonas sp. BIGb0278 | 1 | 3 | 0 | 3 | 0 | 3 | 0 | 1 | 1 | 0 | 0 | 12 |
| Pseudomonas sp. BIGb0408 | 1 | 2 | 1 | 3 | 0 | 2 | 0 | 2 | 1 | 0 | 0 | 12 |
| Pseudomonas sp. MYb115 | 1 | 2 | 0 | 3 | 0 | 4 | 0 | 1 | 1 | 0 | 0 | 12 |
| Pseudomonas sp. MYb187 | 1 | 2 | 0 | 3 | 0 | 4 | 0 | 1 | 1 | 0 | 0 | 12 |
| Pseudomonas sp. MYb60 | 0 | 3 | 0 | 3 | 0 | 3 | 0 | 2 | 1 | 0 | 0 | 12 |
| Pseudomonas trivialis MYb75 | 0 | 3 | 0 | 3 | 0 | 3 | 0 | 2 | 1 | 0 | 0 | 12 |
| Rahnella aquatilis BIGb0112 | 1 | 2 | 0 | 2 | 0 | 1 | 1 | 1 | 3 | 1 | 0 | 12 |
| Rheinheimera pacifica BIGb0367 | 2 | 1 | 0 | 2 | 1 | 1 | 1 | 3 | 1 | 0 | 0 | 12 |
| Achromobacter spanius MYb73 | 0 | 2 | 0 | 2 | 1 | 2 | 2 | 4 | 0 | 0 | 0 | 13 |
| Pseudomonas brenneri BIGb0273 | 0 | 3 | 0 | 3 | 1 | 3 | 0 | 2 | 1 | 0 | 0 | 13 |
| Pseudomonas psychrophila BIGb0222 | 0 | 5 | 0 | 2 | 1 | 3 | 0 | 1 | 1 | 0 | 0 | 13 |
| Pseudomonas putida BIGb0470 | 1 | 3 | 0 | 3 | 0 | 3 | 1 | 1 | 1 | 0 | 0 | 13 |
| Pseudomonas sp. BIGb0381 | 0 | 4 | 0 | 3 | 1 | 3 | 0 | 1 | 1 | 0 | 0 | 13 |
| Pseudomonas sp. JUb96 | 1 | 3 | 0 | 3 | 0 | 4 | 0 | 1 | 1 | 0 | 0 | 13 |
| Pseudomonas sp. MYb2 | 1 | 3 | 0 | 3 | 0 | 4 | 0 | 1 | 1 | 0 | 0 | 13 |
| Pseudomonas sp. MYb3 | 1 | 3 | 0 | 3 | 0 | 4 | 0 | 1 | 1 | 0 | 0 | 13 |

| Genome | COG0376 - Catalase [KatG] | COG0753 - Catalase [KatE] | COG3546 - Mn-containing catalase | COG0386 - Glutathione peroxidase [Gpx] | COG1858 - Cytochrome c peroxidase [Ccp] | COG0450 - Peroxiredoxin [AhpC] | COG1225 - Peroxiredoxin [Bcp] | COG2077 - Peroxiredoxin [Tpx] | COG0678 - Peroxiredoxin [Ahp1] | COG1592 - Rubrerythrin | COG1633 - Rubrerythrin | Total |
|---|---|---|---|---|---|---|---|---|---|---|---|---|
| Rhizobium sp. BIGb0125 | 0 | 3 | 0 | 0 | 4 | 0 | 2 | 2 | 0 | 0 | 2 | 13 |
| Sphingomonas sp. JUb134 | 0 | 3 | 2 | 1 | 0 | 3 | 1 | 3 | 0 | 0 | 0 | 13 |
| Stenotrophomonas rhizophila BIGb0277 | 1 | 2 | 0 | 3 | 2 | 1 | 1 | 3 | 0 | 0 | 0 | 13 |
| Bacillus sp. MYb78 | 0 | 4 | 1 | 1 | 0 | 1 | 0 | 5 | 1 | 0 | 1 | 14 |
| Bacillus subtilis subtilis PS832 | 0 | 3 | 4 | 1 | 0 | 2 | 0 | 2 | 1 | 0 | 1 | 14 |
| Pseudomonas cedrina MYb184 | 0 | 5 | 0 | 3 | 1 | 3 | 0 | 1 | 1 | 0 | 0 | 14 |
| Pseudomonas helmanticensis BIGb0525 | 1 | 3 | 0 | 3 | 1 | 3 | 0 | 2 | 1 | 0 | 0 | 14 |
| Pseudomonas lurida MYb1 | 0 | 3 | 0 | 3 | 0 | 3 | 0 | 4 | 1 | 0 | 0 | 14 |
| Pseudomonas lurida MYb11 | 0 | 3 | 0 | 3 | 0 | 3 | 0 | 4 | 1 | 0 | 0 | 14 |
| Pseudomonas lurida MYb16 | 0 | 3 | 0 | 3 | 0 | 3 | 0 | 4 | 1 | 0 | 0 | 14 |
| Pseudomonas putida BIGb0473 | 1 | 2 | 0 | 3 | 1 | 3 | 0 | 3 | 1 | 0 | 0 | 14 |
| Pseudomonas sp. MYb13 | 0 | 3 | 0 | 3 | 0 | 3 | 0 | 4 | 1 | 0 | 0 | 14 |
| Pseudomonas sp. MYb193 | 0 | 5 | 0 | 3 | 1 | 3 | 0 | 1 | 1 | 0 | 0 | 14 |
| Sphingomonas sp. JUb134 | 0 | 4 | 2 | 1 | 0 | 3 | 1 | 3 | 0 | 0 | 0 | 14 |
| Myroides odoratus BIGb0203 | 0 | 1 | 0 | 1 | 5 | 1 | 0 | 5 | 1 | 1 | 0 | 15 |
| Paenibacillus sp. MYb63 | 0 | 2 | 2 | 4 | 0 | 1 | 0 | 4 | 1 | 0 | 1 | 15 |
| Paenibacillus sp. MYb67 | 0 | 2 | 2 | 4 | 0 | 1 | 0 | 4 | 1 | 0 | 1 | 15 |
| Pseudomonas grimontii BIGb0428 | 0 | 6 | 0 | 3 | 1 | 3 | 0 | 1 | 1 | 0 | 0 | 15 |
| Pseudomonas sp. BIGb0450 | 1 | 4 | 0 | 3 | 1 | 3 | 0 | 2 | 1 | 0 | 0 | 15 |
| Pseudomonas sp. BIGb0558 | 1 | 4 | 0 | 3 | 1 | 3 | 0 | 2 | 1 | 0 | 0 | 15 |
| Pseudomonas poae MYb114 | 0 | 4 | 1 | 3 | 0 | 3 | 0 | 4 | 1 | 0 | 0 | 16 |
| Pseudomonas poae MYb117 | 0 | 4 | 1 | 3 | 0 | 3 | 0 | 4 | 1 | 0 | 0 | 16 |
| Pseudomonas protegens BIGb0404 | 1 | 3 | 0 | 3 | 2 | 3 | 0 | 3 | 1 | 0 | 0 | 16 |
| Pseudomonas protegens JUb28 | 1 | 3 | 0 | 3 | 2 | 3 | 0 | 3 | 1 | 0 | 0 | 16 |
| Pseudomonas sp. BIGb0176 | 1 | 4 | 0 | 3 | 2 | 3 | 0 | 2 | 1 | 0 | 0 | 16 |
| Pseudomonas sp. BIGb0164 | 0 | 7 | 0 | 3 | 0 | 3 | 0 | 3 | 1 | 0 | 0 | 17 |
| Chryseobacterium sp. JUb44 | 1 | 2 | 0 | 2 | 4 | 2 | 0 | 4 | 1 | 2 | 1 | 19 |
| Chryseobacterium sp. MYb7 | 1 | 2 | 0 | 2 | 5 | 2 | 0 | 5 | 2 | 0 | 0 | 19 |
| Pseudomonas sp. JUb42 | 2 | 3 | 2 | 4 | 0 | 3 | 0 | 3 | 1 | 0 | 1 | 19 |
| Chryseobacterium culicis MYb25 | 1 | 2 | 0 | 2 | 6 | 2 | 0 | 6 | 2 | 0 | 0 | 21 |
| Chryseobacterium culicis MYb44 | 1 | 2 | 0 | 2 | 6 | 2 | 0 | 6 | 2 | 0 | 0 | 21 |
| Chryseobacterium nakagawai BIGb0215 | 1 | 1 | 0 | 1 | 8 | 2 | 0 | 6 | 2 | 0 | 0 | 21 |
| Chryseobacterium sp. JUb7 | 1 | 2 | 0 | 1 | 4 | 2 | 0 | 5 | 2 | 2 | 2 | 21 |
| Chryseobacterium sp. BIGb0232 | 1 | 1 | 0 | 1 | 9 | 2 | 0 | 5 | 2 | 2 | 3 | 26 |
| Sphingobacterium sp. BIGb0116 | 1 | 2 | 0 | 3 | 4 | 2 | 0 | 13 | 1 | 0 | 0 | 26 |
| Sphingobacterium sp. JUb56 | 1 | 2 | 0 | 3 | 4 | 2 | 0 | 13 | 1 | 0 | 0 | 26 |
| Sphingobacterium sp. BIGb0165 | 1 | 2 | 0 | 2 | 3 | 2 | 0 | 16 | 1 | 0 | 1 | 28 |
| Sphingobacterium sp. JUb78 | 1 | 3 | 0 | 3 | 6 | 2 | 0 | 12 | 1 | 0 | 0 | 28 |
| Sphingobacterium sp. JUb20 | 1 | 2 | 0 | 3 | 3 | 2 | 0 | 18 | 1 | 0 | 0 | 30 |
| Sphingobacterium sp. JUb21 | 1 | 2 | 0 | 3 | 3 | 2 | 0 | 18 | 1 | 0 | 0 | 30 |
| Sphingobacterium kitahiroshimense JUb78 | 1 | 3 | 0 | 5 | 6 | 2 | 0 | 13 | 1 | 0 | 0 | 31 |

### Supplementary Table 2. Bacterial strains tested for catalase activity.

| Collection | Strain | Genus species | Habitat | Catalase assay: slide test | Catalase assay: quantitative assay (log phase) | Catalase assay: quantitative assay (stationary phase) | Strain source |
|---|---|---|---|---|---|---|---|
| Laboratory | OP50 | E. coli B, uracil auxotroph | | bubbles | n.d. | bubbles (>5 mm) | CGC |
| Laboratory | MG1655 | E. coli K12 F- wild type | | bubbles | bubbles (5 mm/OD600) | bubbles (10 mm) | James Imlay |
| Laboratory | JI377 | E. coli MG1655 ahpCF katG katE | | no bubbles | no bubbles | no bubbles | James Imlay |
| Laboratory | PS832 | Bacillus subtilis Trp+ revertant of strain 168 | | n.d. | bubbles (=<0.5 mm/OD600) | bubbles (1.5 mm/OD600) | Peter Setlow |
| Laboratory | PS2664 | Bacillus subtilis PS832 katA::cat katX::erm Cmr Emr | | n.d. | no bubbles | no bubbles | Peter Setlow |
| CeMbio | MYb10 | Acinetobacter guillouiae | compost | n.d. | n.d. | bubbles (>5 mm/OD600) | CGC (Dirksen et al., 2020) |
| CeMbio | JUb44 | Chryseobacterium scophthalmum | fruit | bubbles | n.d. | bubbles (>5 mm/OD600) | CGC (Dirksen et al., 2020) |
| CeMbio | BIGb0172 | Comamonas piscis | fruit | n.d. | n.d. | bubbles (>5 mm/OD600) | CGC (Dirksen et al., 2020) |
| CeMbio | CEent1 | Enterobacter hormaechei | compost | n.d. | n.d. | bubbles (>5 mm/OD600) | CGC (Dirksen et al., 2020) |
| CeMbio | JUb66 | Lelliottia amnigena | fruit | bubbles | n.d. | bubbles (>5 mm/OD600) | CGC (Dirksen et al., 2020) |
| CeMbio | MYb71 | Ochrobactrum vermis | compost | n.d. | n.d. | bubbles (>5 mm/OD600) | CGC (Dirksen et al., 2020) |
| CeMbio | BIGb0393 | Pantoea nemavictus | fruit | n.d. | n.d. | bubbles (>5 mm/OD600) | CGC (Dirksen et al., 2020) |
| CeMbio | MSPm1 | Pseudomonas berkeleyensis | compost | n.d. | n.d. | bubbles (>5 mm/OD600) | CGC (Dirksen et al., 2020) |
| CeMbio | MYb11 | Pseudomonas lurida | compost | n.d. | n.d. | bubbles (>5 mm/OD600) | CGC (Dirksen et al., 2020) |
| CeMbio | BIGb0170 | Sphingobacterium multivorum | fruit | n.d. | n.d. | bubbles (>5 mm/OD600) | CGC (Dirksen et al., 2020) |
| CeMbio | JUb134 | Sphingomonas molluscorum | fruit | n.d. | n.d. | bubbles (>5 mm/OD600) | CGC (Dirksen et al., 2020) |
| CeMbio | JUb19 | Stenotrophomonas indicatrix | fruit | bubbles | n.d. | bubbles (>5 mm/OD600) | CGC (Dirksen et al., 2020) |
| BIGb | BIGb0220 | Lactococcus lactis | fruit | no bubbles | n.d. | n.d. | Buck Samuel (Samuel et. al 2016) |
| JUb | JUb24 | Achromobacter | fruit | bubbles | n.d. | n.d. | Gary Ruvkun (Samuel et. al 2016) |
| JUb | JUb104 | Achromobacter | snail or slug | bubbles | n.d. | n.d. | Gary Ruvkun (Samuel et. al 2016) |
| JUb | JUb89 | Acinetobacter | snail or slug | bubbles | n.d. | n.d. | Gary Ruvkun (Samuel et. al 2016) |
| JUb | JUb2 | Acinetobacter | compost | bubbles | n.d. | n.d. | Gary Ruvkun (Samuel et. al 2016) |
| JUb | JUb81 | Acinetobacter | fruit | bubbles | n.d. | n.d. | Gary Ruvkun (Samuel et. al 2016) |
| JUb | JUb68 | Acinetobacter | fruit | bubbles | n.d. | n.d. | Gary Ruvkun (Samuel et. al 2016) |
| JUb | JUb108 | Acinetobacter | snail or slug | bubbles | n.d. | n.d. | Gary Ruvkun (Samuel et. al 2016) |
| JUb | JUb109 | Acinetobacter | snail or slug | bubbles | n.d. | n.d. | Gary Ruvkun (Samuel et. al 2016) |
| JUb | JUb118 | Acinetobacter | snail or slug | bubbles | n.d. | n.d. | Gary Ruvkun (Samuel et. al 2016) |
| JUb | JUb110 | Acinetobacter | snail or slug | bubbles | n.d. | n.d. | Gary Ruvkun (Samuel et. al 2016) |
| JUb | JUb4 | Alcaligenes | compost | bubbles | n.d. | n.d. | Gary Ruvkun (Samuel et. al 2016) |
| JUb | JUb11 | Bacillus | compost | bubbles | n.d. | n.d. | Gary Ruvkun (Samuel et. al 2016) |
| JUb | JUb14 | Bacillus | compost | bubbles | n.d. | n.d. | Gary Ruvkun (Samuel et. al 2016) |
| JUb | JUb113 | Bacillus | snail or slug | bubbles | n.d. | n.d. | Gary Ruvkun (Samuel et. al 2016) |
| JUb | JUb91 | Bacillus | snail or slug | bubbles | n.d. | n.d. | Gary Ruvkun (Samuel et. al 2016) |
| JUb | JUb32 | Bacillus | fruit | bubbles | n.d. | n.d. | Gary Ruvkun (Samuel et. al 2016) |
| JUb | JUb87 | Buttiauxella | snail or slug | bubbles | n.d. | n.d. | Gary Ruvkun (Samuel et. al 2016) |
| JUb | JUb37 | Cellulomonas | fruit | bubbles | n.d. | n.d. | Gary Ruvkun (Samuel et. al 2016) |
| JUb | JUb7 | Chryseobacterium | compost | bubbles | n.d. | n.d. | Gary Ruvkun (Samuel et. al 2016) |
| JUb | JUb58 | Comamonas | fruit | bubbles | n.d. | n.d. | Gary Ruvkun (Samuel et. al 2016) |
| JUb | JUb34 | Curtobacterium | fruit | bubbles | n.d. | n.d. | Gary Ruvkun (Samuel et. al 2016) |
| JUb | JUb36 | Curtobacterium | fruit | bubbles | n.d. | n.d. | Gary Ruvkun (Samuel et. al 2016) |
| JUb | JUb8 | Delftia | compost | bubbles | n.d. | n.d. | Gary Ruvkun (Samuel et. al 2016) |
| JUb | JUb105 | Enterobacter | snail or slug | bubbles | n.d. | n.d. | Gary Ruvkun (Samuel et. al 2016) |
| JUb | JUb101 | Enterobacter | snail or slug | bubbles | n.d. | n.d. | Gary Ruvkun (Samuel et. al 2016) |
| JUb | JUb103 | Enterobacter | snail or slug | bubbles | n.d. | n.d. | Gary Ruvkun (Samuel et. al 2016) |
| JUb | JUb25 | Erwinia | fruit | bubbles | n.d. | n.d. | Gary Ruvkun (Samuel et. al 2016) |
| JUb | JUb29 | Erwinia | fruit | bubbles | n.d. | n.d. | Gary Ruvkun (Samuel et. al 2016) |
| JUb | JUb43 | Flavobacterium | fruit | bubbles | n.d. | n.d. | Gary Ruvkun (Samuel et. al 2016) |
| JUb | JUb59 | Lactococcus | fruit | bubbles | n.d. | n.d. | Gary Ruvkun (Samuel et. al 2016) |
| JUb | JUb67 | Lactococcus | fruit | bubbles | n.d. | bubbles (>5 mm/OD600) | Gary Ruvkun (Samuel et. al 2016) |
| JUb | JUb97 | Lactococcus | snail or slug | bubbles | n.d. | bubbles (>5 mm/OD600) | Gary Ruvkun (Samuel et. al 2016) |
| JUb | JUb18 | Leucobacter | lab isolate? | bubbles | n.d. | n.d. | Gary Ruvkun (Samuel et. al 2016) |
| JUb | JUb74 | Microbacterium | fruit | bubbles | n.d. | n.d. | Gary Ruvkun (Samuel et. al 2016) |
| JUb | JUb90 | Microbacterium | snail or slug | bubbles | n.d. | n.d. | Gary Ruvkun (Samuel et. al 2016) |
| JUb | JUb75 | Microbacterium | fruit | bubbles | n.d. | n.d. | Gary Ruvkun (Samuel et. al 2016) |
| JUb | JUb76 | Microbacterium | fruit | bubbles | n.d. | n.d. | Buck Samuel (Samuel et. al 2016) |
| JUb | JUb76b * | Microbacterium | fruit | very few bubles | no bubbles | bubbles (=<0.5 mm/OD600) | Gary Ruvkun (Samuel et. al 2016) |
| JUb | JUb122 | Micrococcus | fruit | bubbles | n.d. | n.d. | Gary Ruvkun (Samuel et. al 2016) |
| JUb | JUb30 | Pantoea | fruit | bubbles | n.d. | n.d. | Gary Ruvkun (Samuel et. al 2016) |
| JUb | JUb5 | Providencia | compost | bubbles | n.d. | n.d. | Gary Ruvkun (Samuel et. al 2016) |
| JUb | JUb39 | Providencia | fruit | bubbles | n.d. | bubbles (>5 mm/OD600) | Gary Ruvkun (Samuel et. al 2016) |
| JUb | JUb102 | Providencia | snail or slug | bubbles | n.d. | n.d. | Gary Ruvkun (Samuel et. al 2016) |
| JUb | JUb6 | Pseudomonas | compost | bubbles | n.d. | n.d. | Gary Ruvkun (Samuel et. al 2016) |
| JUb | JUb85 | Pseudomonas | snail or slug | bubbles | n.d. | n.d. | Gary Ruvkun (Samuel et. al 2016) |
| JUb | JUb114 | Pseudomonas | snail or slug | bubbles | n.d. | n.d. | Gary Ruvkun (Samuel et. al 2016) |
| JUb | JUb95 | Pseudomonas | snail or slug | bubbles | n.d. | n.d. | Gary Ruvkun (Samuel et. al 2016) |
| JUb | JUb100 | Pseudomonas | snail or slug | bubbles | n.d. | n.d. | Gary Ruvkun (Samuel et. al 2016) |
| JUb | JUb12 | Pseudomonas | compost | bubbles | n.d. | n.d. | Gary Ruvkun (Samuel et. al 2016) |
| JUb | JUb42 | Pseudomonas | fruit | bubbles | n.d. | n.d. | Gary Ruvkun (Samuel et. al 2016) |
| JUb | JUb52 | Pseudomonas | fruit | bubbles | n.d. | n.d. | Gary Ruvkun (Samuel et. al 2016) |
| JUb | JUb93 | Pseudomonas putida | snail or slug | bubbles | n.d. | n.d. | Gary Ruvkun (Samuel et. al 2016) |
| JUb | JUb38 | Raoultella | fruit | bubbles | n.d. | n.d. | Gary Ruvkun (Samuel et. al 2016) |
| JUb | JUb54 | Raoultella | fruit | bubbles | n.d. | n.d. | Gary Ruvkun (Samuel et. al 2016) |
| JUb | JUb99 | Raoultella | snail or slug | bubbles | n.d. | n.d. | Gary Ruvkun (Samuel et. al 2016) |
| JUb | JUb83 | Rhodococcus | fruit | bubbles | n.d. | n.d. | Gary Ruvkun (Samuel et. al 2016) |
| JUb | JUb9 | Serratia | compost | bubbles | n.d. | n.d. | Gary Ruvkun (Samuel et. al 2016) |
| JUb | JUb20 | Sphingobacterium | fruit | bubbles | n.d. | n.d. | Gary Ruvkun (Samuel et. al 2016) |
| JUb | JUb21 | Sphingobacterium | fruit | bubbles | n.d. | n.d. | Gary Ruvkun (Samuel et. al 2016) |
| JUb | JUb79 | Sphingobacterium | fruit | bubbles | n.d. | n.d. | Gary Ruvkun (Samuel et. al 2016) |
| JUb | JUb56 | Sphingobacterium | fruit | bubbles | n.d. | n.d. | Gary Ruvkun (Samuel et. al 2016) |
| JUb | JUb86 | Sphingobacterium | snail or slug | bubbles | n.d. | n.d. | Gary Ruvkun (Samuel et. al 2016) |
| JUb | JUb94 | Sphingobacterium | snail or slug | bubbles | n.d. | n.d. | Gary Ruvkun (Samuel et. al 2016) |
| JUb | JUb78 | Sphingobacterium | fruit | bubbles | n.d. | n.d. | Gary Ruvkun (Samuel et. al 2016) |
| JUb | JUb15 | Staphylococcus | fruit | bubbles | n.d. | n.d. | Gary Ruvkun (Samuel et. al 2016) |
| JUb | JUb23 | Stenotrophomonas | fruit | bubbles | n.d. | n.d. | Gary Ruvkun (Samuel et. al 2016) |
| MOYb | MOYb_037 | Acinetobacter | compost | bubbles | n.d. | n.d. | This study |
| MOYb | MOYb_079 | Acinetobacter | compost | bubbles | n.d. | n.d. | This study |
| MOYb | MOYb_081 | Acinetobacter | compost | bubbles | n.d. | n.d. | This study |
| MOYb | MOYb_092 | Acinetobacter | compost | bubbles | n.d. | n.d. | This study |
| MOYb | MOYb_032 | Acinetobacter johnsonii | compost | bubbles | n.d. | n.d. | This study |
| MOYb | MOYb_080 | Acintetobacter | compost | bubbles | n.d. | n.d. | This study |
| MOYb | MOYb_027 | Arthrobacter | compost | bubbles | n.d. | n.d. | This study |
| MOYb | MOYb_029 | Arthrobacter | compost | bubbles | n.d. | n.d. | This study |
| MOYb | MOYb_044 | Arthrobacter | compost | bubbles | n.d. | n.d. | This study |
| MOYb | MOYb_045 | Arthrobacter | compost | bubbles | n.d. | n.d. | This study |
| MOYb | MOYb_052 | Arthrobacter | compost | bubbles | n.d. | n.d. | This study |
| MOYb | MOYb_054 | Arthrobacter | compost | bubbles | n.d. | n.d. | This study |
| MOYb | MOYb_064 | Arthrobacter | compost | bubbles | n.d. | n.d. | This study |
| MOYb | MOYb_077 | Arthrobacter | compost | bubbles | n.d. | n.d. | This study |
| MOYb | MOYb_082 | Arthrobacter | compost | bubbles | n.d. | n.d. | This study |
| MOYb | MOYb_066 | Arthrobacter | compost | bubbles | n.d. | n.d. | This study |
| MOYb | MOYb_034 | Athrobacter | compost | bubbles | n.d. | n.d. | This study |

| Collection | Strain | Genus species | Habitat | Catalase assay: slide test | Catalase assay: quantitative assay (log phase) | Catalase assay: quantitative assay (stationary phase) | Strain source |
|---|---|---|---|---|---|---|---|
| MOYb | MOYb_042 | Bacillus | compost | bubbles | n.d. | n.d. | This study |
| MOYb | MOYb_048 | Bacillus | compost | bubbles | n.d. | n.d. | This study |
| MOYb | MOYb_050 | Bacillus | compost | bubbles | n.d. | n.d. | This study |
| MOYb | MOYb_051 | Bacillus | compost | bubbles | n.d. | n.d. | This study |
| MOYb | MOYb_053 | Bacillus | compost | bubbles | n.d. | n.d. | This study |
| MOYb | MOYb_088 | Bacillus | compost | bubbles | n.d. | n.d. | This study |
| MOYb | MOYb_089 | Bacillus | compost | bubbles | n.d. | n.d. | This study |
| MOYb | MOYb_095 | Bacillus | compost | bubbles | n.d. | n.d. | This study |
| MOYb | MOYb_098 | Bacillus | compost | bubbles | n.d. | n.d. | This study |
| MOYb | MOYb_100 | Bacillus | compost | bubbles | n.d. | n.d. | This study |
| MOYb | MOYb_105 | Bacillus | compost | bubbles | n.d. | n.d. | This study |
| MOYb | MOYb_083 | Bacillus pumilus | compost | bubbles | n.d. | n.d. | This study |
| MOYb | MOYb_102 | Citrobacter | compost | bubbles | n.d. | n.d. | This study |
| MOYb | MOYb_101 | Enterobacter | compost | bubbles | n.d. | n.d. | This study |
| MOYb | MOYb_106 | Enterobacter/Klebsiella | compost | bubbles | n.d. | n.d. | This study |
| MOYb | MOYb_109 | Enterobacter/Klebsiella | compost | bubbles | n.d. | n.d. | This study |
| MOYb | MOYb_084 | Erwinia or Pantoea | compost | bubbles | n.d. | n.d. | This study |
| MOYb | MOYb_094 | Hafnia | compost | bubbles | n.d. | n.d. | This study |
| MOYb | MOYb_055 | Klebsiella | compost | bubbles | n.d. | n.d. | This study |
| MOYb | MOYb_060 | Klebsiella | compost | bubbles | n.d. | n.d. | This study |
| MOYb | MOYb_061 | Klebsiella | compost | bubbles | n.d. | n.d. | This study |
| MOYb | MOYb_086 | Microbacterium | compost | bubbles | n.d. | n.d. | This study |
| MOYb | MOYb_087 | Microbacterium | compost | bubbles | n.d. | n.d. | This study |
| MOYb | MOYb_085 | Paenibacillus | compost | bubbles | n.d. | n.d. | This study |
| MOYb | MOYb_057 | Paracoccus | compost | bubbles | n.d. | n.d. | This study |
| MOYb | MOYb_090 | Phyllobacteriaceae | compost | bubbles | n.d. | n.d. | This study |
| MOYb | MOYb_023 | Pseudomonas | compost | bubbles | n.d. | n.d. | This study |
| MOYb | MOYb_024 | Pseudomonas | compost | bubbles | n.d. | n.d. | This study |
| MOYb | MOYb_026 | Pseudomonas | compost | bubbles | n.d. | n.d. | This study |
| MOYb | MOYb_031 | Pseudomonas | compost | bubbles | n.d. | n.d. | This study |
| MOYb | MOYb_036 | Pseudomonas | compost | bubbles | n.d. | n.d. | This study |
| MOYb | MOYb_038 | Pseudomonas | compost | bubbles | n.d. | n.d. | This study |
| MOYb | MOYb_039 | Pseudomonas | compost | bubbles | n.d. | n.d. | This study |
| MOYb | MOYb_040 | Pseudomonas | compost | bubbles | n.d. | n.d. | This study |
| MOYb | MOYb_041 | Pseudomonas | compost | bubbles | n.d. | n.d. | This study |
| MOYb | MOYb_043 | Pseudomonas | compost | bubbles | n.d. | n.d. | This study |
| MOYb | MOYb_046 | Pseudomonas | compost | bubbles | n.d. | n.d. | This study |
| MOYb | MOYb_049 | Pseudomonas | compost | bubbles | n.d. | n.d. | This study |
| MOYb | MOYb_056 | Pseudomonas | compost | bubbles | n.d. | n.d. | This study |
| MOYb | MOYb_059 | Pseudomonas | compost | bubbles | n.d. | n.d. | This study |
| MOYb | MOYb_062 | Pseudomonas | compost | bubbles | n.d. | n.d. | This study |
| MOYb | MOYb_063 | Pseudomonas | compost | bubbles | n.d. | n.d. | This study |
| MOYb | MOYb_075 | Pseudomonas | compost | bubbles | n.d. | n.d. | This study |
| MOYb | MOYb_078 | Pseudomonas | compost | bubbles | n.d. | n.d. | This study |
| MOYb | MOYb_091 | Pseudomonas | compost | bubbles | n.d. | n.d. | This study |
| MOYb | MOYb_033 | Pseudomonas kilonensis | compost | bubbles | n.d. | n.d. | This study |
| MOYb | MOYb_107 | Rhodotorula yeast | compost | bubbles | n.d. | n.d. | This study |
| MOYb | MOYb_108 | Rhodotorula yeast | compost | bubbles | n.d. | n.d. | This study |
| MOYb | MOYb_093 | Serratia | compost | bubbles | n.d. | n.d. | This study |
| MOYb | MOYb_096 | Serratia | compost | bubbles | n.d. | n.d. | This study |
| MOYb | MOYb_097 | Serratia | compost | bubbles | n.d. | n.d. | This study |
| MOYb | MOYb_099 | Serratia | compost | bubbles | n.d. | n.d. | This study |
| MOYb | MOYb_021 | Shewanella | compost | very few bubles | no bubbles | no bubbles | This study |
| MOYb | MOYb_025 | Shewanella | compost | very few bubles | no bubbles | no bubbles | This study |
| MOYb | MOYb_028 | Shewanella | compost | very few bubles | no bubbles | no bubbles | This study |
| MOYb | MOYb_035 | Shewanella | compost | very few bubles | no bubbles | no bubbles | This study |
| MOYb | MOYb_058 | Shewanella | compost | very few bubles | no bubbles | no bubbles | This study |
| MOYb | MOYb_067 | Shewanella | compost | very few bubles | no bubbles | no bubbles | This study |
| MOYb | MOYb_068 | Shewanella | compost | very few bubles | no bubbles | no bubbles | This study |
| MOYb | MOYb_069 | Shewanella | compost | very few bubles | no bubbles | no bubbles | This study |
| MOYb | MOYb_070 | Shewanella | compost | very few bubles | no bubbles | no bubbles | This study |
| MOYb | MOYb_071 | Shewanella | compost | very few bubles | no bubbles | no bubbles | This study |
| MOYb | MOYb_072 | Shewanella | compost | very few bubles | no bubbles | no bubbles | This study |
| MOYb | MOYb_073 | Shewanella | compost | very few bubles | no bubbles | no bubbles | This study |
| MOYb | MOYb_074 | Shewanella | compost | very few bubles | no bubbles | no bubbles | This study |
| MOYb | MOYb_076 | Shewanella | compost | very few bubles | no bubbles | no bubbles | This study |
| MOYb | MOYb_065 | Trichosporonaceae yeast | compost | bubbles | n.d. | n.d. | This study |
| MOYb | MOYb_103 | | compost | bubbles | n.d. | n.d. | This study |
| MOYb | MOYb_104 | | compost | bubbles | n.d. | n.d. | This study |
| MOYb | MOYb_022 | | compost | bubbles | n.d. | n.d. | This study |

* The JUb76 strain isolate from Buck Samuel's lab was catalase positive, but the JUb76 strain isolate from Gary Ruvkun's lab that we originally screened was catalase negative. We refer to this latter strain as JUb76b, since its partial 16S rRNA gene sequence matched the respective published sequenced for JUb76 (Samuel et. al 2016).

n.d. = assay not done

#### Supplementary Table 3. Bacterial strains.

| Strain | Genotype | Source | Reference |
|---|---|---|---|
| OP50 | *E. coli* B, uracil auxotroph | CGC | Brenner, 1974 |
| HT115 | *E. coli* $F^-$ *mcrA, mcrB, IN(rrnD-rrnE)1, rnc14::Tn10(DE3 lysogen: lavUV5 promoter -T7 polymerase).* Tetracycline resistant | CGC | Kamath et al., 2001 |
| MG1655 | *E. coli* K12 $F^-$ wild type | James Imlay | Seaver and Imlay, 2001 |
| JI377 | *E. coli* MG1655 *ahpCF katG katE* | James Imlay | Seaver and Imlay, 2001 |
| PS832 | *Bacillus subtilis* $Trp^+$ revertant of strain 168 | Peter Setlow | Bagyan et al., 1998 |
| PS2664 | *Bacillus subtilis* PS832 *katA::cat katX::erm* $Cm^r$ $Em^r$ | Peter Setlow | Bagyan et al., 1998 |
| MYb71 | *Ochrobactrum vermis* | CGC | Dirksen et al., 2016 |
| JUb39 | *Providencia alcalifaciens* | Piali Sengupta | Samuel et al., 2016 |
| JUb45 | *Neorhizobium huautlense* | Gary Ruvkun | Samuel et al., 2016 |

#### Supplementary Table 4. Nematode strains.

| Strain | Species | Genotype | Source | Reference |
|---|---|---|---|---|
| QZ0 | *Caenorhabditis elegans* | N2 wild type | Joy Alcedo | Schiffer et. al, 2020 |
| AF16 | *Caenorhabditis briggsae* | wild isolate from soil in Ahmedabad, India | CGC | |
| JU322 | *Caenorhabditis elegans* | wild isolate from a Helix snail on the trunk of a mulberry tree in Merlet, Lagorce (Ardeche), France | CGC | Barriere and Felix, 2005 |
| PX178 | *Caenorhabditis elegans* | wild isolate from a snail under rocks over damp organic soil in northwest Hendricks Park, Eugene, OR (USA) | CGC | |
| MT14984 | *Caenorhabditis elegans* | *tph-1(n4622)* | CGC | Shivers et al., 2009 |
| MT15434 | *Caenorhabditis elegans* | *tph-1(mg280)* . In this backcrossed *mg280* allele a *cam-1* mutation was removed by crossing left and right of *tph-1(mg280)* using *bli-2* and *unc-4* . Strain does not have withered tail defect and moves well. | CGC | Sze et al., 2000 |
| MT8944 | *Caenorhabditis elegans* | *mod-5(n822)* | CGC | Ranganathan et al., 2001 |
| ZD763 | *Caenorhabditis elegans* | *mgIs40[Pdaf-28::nls-GFP]; jxEx102[Ptrx-1::ICE + Pofm-1::gfp]* | Dennis Kim | Cornils et al., 2011 |
| ZM10104 | *Caenorhabditis elegans* | *Pift-20::GCaMP6::3xNLS + Pgpc-1::mCherry* | This work | |

### Supplementary Table 5. MOYb nematode microbiome collection from residential compost.

| MOYb_strain | Genus/species | Isolation notes | Associated worm | 16S rRNA gene primers | 16S rRNA gene sequence (if available) | Additional notes |
|---|---|---|---|---|---|---|
| MOYb_001 | tbd | compost | | | | |
| MOYb_002 | tbd | compost | | | | |
| MOYb_003 | tbd | compost | | | | |
| MOYb_004 | Micrococcus luteus | lab isolate | | 27f/1492r | | |
| MOYb_005 | Arthrobacter sp. | lab isolate | | 27f/1492r | | |
| MOYb_006 | tbd | Tomato#1_A, e.coli-like odor | MO_w003,MO_w004 | | | |
| MOYb_007 | Providencia rettgeri | Tomato#1_B, nutty odor | MO_w003,MO_w004 | | | |
| MOYb_008 | Klebsiella sp. (oxytoca/pneumonia) | Tomato#1_C1 | MO_w003,MO_w004 | 27f/1492r | | |
| MOYb_009 | Chryseobacterium sp. (lathyri) | Tomato#1_C2 | MO_w003,MO_w004 | 515f/806rB | | |
| MOYb_010 | Pseudomonas sp. | Tomato#1_D | MO_w003,MO_w004 | 515f/806rB | | worms dead |
| MOYb_011 | Chryseobacterium sp. (indologenes) - weak homology | Tomato#1_E, fruity odor | MO_w003,MO_w004 | 515f/806rB | | |
| MOYb_012 | Rahnella sp. - questionable | Tomato#1_F | MO_w003,MO_w004 | 27f/1492r | | |
| MOYb_013 | Stenotrophomonas/Microbacterium sp. | Tomato#2_A1 | MO_w005 | 515f/806rB | same sequence match for Stenotrophomonas, Rhizobium sp. | |
| MOYb_014 | Myroides sp. (odoratimimus)? | Tomato#2_A2 | MO_w005 | 515f/806rB | | |
| MOYb_015 | tbd | Tomato#2_B | MO_w005 | | | |
| MOYb_016 | tbd | Tomato#2_C | MO_w005 | | | |
| MOYb_017 | Pseudomonas sp. - unclear | Tomato#2_D | MO_w005 | 515f/806rB | | |
| MOYb_018 | Klebsiella sp. (oxytoca) | Dirt#2_A, fruity odor | MO_w006 | 27f/1492r | | |
| MOYb_019 | Microbacterium sp. | Dirt#2_B1 | MO_w006 | 515f/806rB | | worms dead |
| MOYb_020 | E. coli | Dirt#2_B2 | MO_w006 | 27f/1492r | | |
| MOYb_021 | Shewanella | JU1_1 | JU322_1 wormpost | 515f/806rB | [illegible] | |
| MOYb_022 | | | | | | |
| MOYb_023 | Pseudomonas | JU1_3 | JU322_1 wormpost | 515f/806rB | [illegible] | |
| MOYb_024 | Pseudomonas | JU1_4 | JU322_1 wormpost | 515f/806rB | [illegible] | |
| MOYb_025 | Shewanella | JU1_5 | JU322_1 wormpost | 515f/806rB | [illegible] | |
| MOYb_026 | Pseudomonas | JU1_6 | JU322_1 wormpost | 515f/806rB | [illegible] | |
| MOYb_027 | Arthrobacter | JU1_7 | JU322_1 wormpost | 515f/806rB | [illegible] | |
| MOYb_028 | Shewanella | JU1_8 | JU322_1 wormpost | 515f/806rB | [illegible] | |
| MOYb_029 | Arthrobacter | JU1_9 | JU322_1 wormpost | 515f/806rB | [illegible] | |
| MOYb_030 | Bacillus cereus | JU1_10 | JU322_1 wormpost | 515f/806rB | [illegible] | |
| MOYb_031 | Pseudomonas | JU1_11 | JU322_1 wormpost | 515f/806rB | [illegible] | |
| MOYb_032 | Acinetobacter johnsonii | JU1_12 | JU322_1 wormpost | 515f/806rB | [illegible] | |
| MOYb_033 | Pseudomonas kilonensis | JU1_13 | JU322_1 wormpost | 515f/806rB | [illegible] | |
| MOYb_034 | Athrobacter | JU1_14 | JU322_1 wormpost | 515f/806rB | [illegible] | |
| MOYb_035 | Shewanella | JU1_15 | JU322_1 wormpost | 515f/806rB | [illegible] | |
| MOYb_036 | Pseudomonas | JU1_16 | JU322_1 wormpost | 515f/806rB | [illegible] | |
| MOYb_037 | Acinetobacter | JU1_17 | JU322_1 wormpost | 515f/806rB | [illegible] | |
| MOYb_038 | Pseudomonas | JU1_18 | JU322_1 wormpost | 515f/806rB | [illegible] | |
| MOYb_039 | Pseudomonas | JU2_19 | JU322_2 wormpost | 515f/806rB | [illegible] | |
| MOYb_040 | Pseudomonas | JU2_20 | JU322_2 wormpost | 515f/806rB | [illegible] | |
| MOYb_041 | Pseudomonas | JU2_21 | JU322_2 wormpost | 515f/806rB | [illegible] | |
| MOYb_042 | Bacillus | JU2_22 | JU322_2 wormpost | 515f/806rB | [illegible] | |
| MOYb_043 | Pseudomonas | JU2_23 | JU322_2 wormpost | 515f/806rB | [illegible] | |
| MOYb_044 | Arthrobacter | JU2_24 | JU322_2 wormpost | 515f/806rB | [illegible] | |
| MOYb_045 | Arthrobacter | JU2_25 | JU322_2 wormpost | 515f/806rB | [illegible] | |
| MOYb_046 | Pseudomonas | JU2_26 | JU322_2 wormpost | 515f/806rB | [illegible] | |
| MOYb_047 | Bacillus | JU2_27 | JU322_2 wormpost | 515f/806rB | [illegible] | |
| MOYb_048 | Bacillus | JU3_28 | JU322_3 wormpost | 515f/806rB | [illegible] | |
| MOYb_049 | Pseudomonas | JU3_29 | JU322_3 wormpost | 515f/806rB | [illegible] | |
| MOYb_050 | Bacillus | JU3_30 | JU322_3 wormpost | 515f/806rB | [illegible] | |
| MOYb_051 | Bacillus | N2_3_1 | N2_3_wormpost | 515f/806rB | [illegible] | |
| MOYb_052 | Arthrobacter | N2_3_2 | N2_3_wormpost | 515f/806rB | [illegible] | |
| MOYb_053 | Bacillus | N2_3_3 | N2_3_wormpost | 515f/806rB | [illegible] | |
| MOYb_054 | Arthrobacter | N2_3_4 | N2_3_wormpost | 515f/806rB | [illegible] | |
| MOYb_055 | Klebsiella | N2_3_5 | N2_3_wormpost | 515f/806rB | [illegible] | |
| MOYb_056 | Pseudomonas | N2_3_6 | N2_3_wormpost | 515f/806rB | [illegible] | |
| MOYb_057 | Paracoccus | N2_3_7 | N2_3_wormpost | 515f/806rB | [illegible] | |
| MOYb_058 | Shewanella | N2_3_8 | N2_3_wormpost | 515f/806rB | [illegible] | |
| MOYb_059 | Pseudomonas | N2_3_9 | N2_3_wormpost | 515f/806rB | [illegible] | |
| MOYb_060 | Klebsiella | N2_3_10 | N2_3_wormpost | 515f/806rB | [illegible] | |
| MOYb_061 | Klebsiella | N2_3_11 | N2_3_wormpost | 515f/806rB | [illegible] | |
| MOYb_062 | Pseudomonas | N2_3_12 | N2_3_wormpost | 515f/806rB | [illegible] | |
| MOYb_063 | Pseudomonas | N2_3_13 | N2_3_wormpost | 515f/806rB | [illegible] | |
| MOYb_064 | Arthrobacter | N2_3_14 | N2_3_wormpost | 515f/806rB | [illegible] | |
| MOYb_065 | Trichosporonaceae yeast | N2_3_15 | N2_3_wormpost | 515f/806rB | [illegible] | |
| MOYb_066 | Arthrobcter | N2_3_16 | N2_3_wormpost | 515f/806rB | [illegible] | |
| MOYb_067 | Shewanella | N2_3_17 | N2_3_wormpost | 515f/806rB | [illegible] | |
| MOYb_068 | Shewanella | N2_3_18 | N2_3_wormpost | 515f/806rB | [illegible] | |
| MOYb_069 | Shewanella | N2_3_19 | N2_3_wormpost | 515f/806rB | [illegible] | |
| MOYb_070 | Shewanella | N2_3_20 | N2_3_wormpost | 515f/806rB | [illegible] | |
| MOYb_071 | Shewanella | N2_2_21 | N2_2_wormpost | 515f/806rB | [illegible] | |
| MOYb_072 | Shewanella | N2_2_22 | N2_2_wormpost | 515f/806rB | [illegible] | |
| MOYb_073 | Shewanella | N2_2_23 | N2_2_wormpost | 515f/806rB | [illegible] | |
| MOYb_074 | Shewanella | N2_2_24 | N2_2_wormpost | 515f/806rB | [illegible] | |
| MOYb_075 | Pseudomonas | N2_2_25 | N2_2_wormpost | 515f/806rB | [illegible] | |
| MOYb_076 | Shewanella | N2_2_26 | N2_2_wormpost | 515f/806rB | [illegible] | |
| MOYb_077 | Arthrobacter | N2_2_27 | N2_2_wormpost | 515f/806rB | [illegible] | |
| MOYb_078 | Pseudomonas | N2_1_28 | N2_1_wormpost | 515f/806rB | [illegible] | |
| MOYb_079 | Acinetobacter | N2_1_29 | N2_1_wormpost | 515f/806rB | [illegible] | |
| MOYb_080 | Acintetobacter | N2_1_30 | N2_1_wormpost | 515f/806rB | [illegible] | |
| MOYb_081 | Acinetobacter | N2_1_31 | N2_1_wormpost | 515f/806rB | [illegible] | |
| MOYb_082 | Arthrobacter | N2_1_32 | N2_1_wormpost | 515f/806rB | [illegible] | |
| MOYb_083 | Bacillus pumilus | PX1_1 | PX178_1_wormpost | 515f/806rB | [illegible] | |
| MOYb_084 | Erwinia or Pantoea | PX1_2 | PX178_1_wormpost | 515f/806rB | [illegible] | |
| MOYb_085 | Paenibacillus | PX1_3 | PX178_1_wormpost | 515f/806rB | [illegible] | |
| MOYb_086 | Microbacterium | PX1_4 | PX178_1_wormpost | 515f/806rB | [illegible] | |
| MOYb_087 | Microbacterium | PX1_5 | PX178_1_wormpost | 515f/806rB | [illegible] | |
| MOYb_088 | Bacillus | PX2_6 | PX178_2_wormpost | 515f/806rB | [illegible] | |
| MOYb_089 | Bacillus | PX2_7 | PX178_2_wormpost | 515f/806rB | [illegible] | |
| MOYb_090 | Phyllobacteriaceae | PX2_8 | PX178_2_wormpost | 515f/806rB | [illegible] | |
| MOYb_091 | Pseudomonas | PX2_9 | PX178_2_wormpost | 515f/806rB | [illegible] | |
| MOYb_092 | Acinetobacter | PX2_10 | PX178_2_wormpost | 515f/806rB | [illegible] | |
| MOYb_093 | Serratia | PX3_11 | PX178_3_wormpost | 515f/806rB | [illegible] | |
| MOYb_094 | Hafnia | PX3_12 | PX178_3_wormpost | 515f/806rB | [illegible] | |
| MOYb_095 | Bacillus | PX3_13 | PX178_3_wormpost | 515f/806rB | [illegible] | |
| MOYb_096 | Serratia | PX3_14 | PX178_3_wormpost | 515f/806rB | [illegible] | |
| MOYb_097 | Serratia | PX3_15 | PX178_3_wormpost | 515f/806rB | [illegible] | |
| MOYb_098 | Bacillus | PX3_16 | PX178_3_wormpost | 515f/806rB | [illegible] | |
| MOYb_099 | Serratia | PX3_17 | PX178_3_wormpost | 515f/806rB | [illegible] | |
| MOYb_100 | Bacillus | PX3_18 | PX178_3_wormpost | 515f/806rB | [illegible] | |
| MOYb_101 | Enterobacter | PX3_19 | PX178_3_wormpost | 515f/806rB | [illegible] | |
| MOYb_102 | Citrobacter | PX3_20 | PX178_3_wormpost | 515f/806rB | [illegible] | |
| MOYb_103 | NA | PX3_21 | PX178_3_wormpost | 515f/806rB | NA | |
| MOYb_104 | NA | PX3_22 | PX178_3_wormpost | 515f/806rB | NA | |
| MOYb_105 | Bacillus | PX3_23 | PX178_3_wormpost | 515f/806rB | [illegible] | |
| MOYb_106 | Enterobacter/Klebsiella | PX3_24 | PX178_3_wormpost | 515f/806rB | [illegible] | |
| MOYb_107 | Rhodotorula yeast | PX3_25 | PX178_3_wormpost | 515f/806rB | [illegible] | |
| MOYb_108 | Rhodotorula yeast | PX3_26 | PX178_3_wormpost | 515f/806rB | [illegible] | |
| MOYb_109 | Enterobacter/Klebsiella | PX3_27 | PX178_3_wormpost | 515f/806rB | [illegible] | |
